# Climate change should drive mammal defaunation in tropical dry forests

**DOI:** 10.1101/2023.08.17.553094

**Authors:** Mario R. Moura, Gibran A. Oliveira, Adriano P. Paglia, Mathias M. Pires, Bráulio A. Santos

## Abstract

Human-induced climate change has intensified negative impacts on socioeconomic factors, the environment, and biodiversity, including changes in rainfall patterns and an increase in global average temperatures. Drylands are particularly at risk, with projections suggesting they will become hotter, drier, and less suitable for a significant portion of their species, potentially leading to mammal defaunation. We use ecological niche modelling and community ecology biodiversity metrics to examine potential geographical range shifts of non-volant mammal species in the largest Neotropical dryland, the Caatinga, and evaluate impacts of climate change on mammal assemblages. According to projections, 85% of the mammal species will lose suitable habitats, with one quarter of species projected to completely lose suitable habitats by 2060. This will result in a decrease in species richness for more than 90% of assemblages and an increase in compositional similarity to nearby assemblages (i.e., reduction in spatial beta diversity) for 70% of the assemblages. Small-sized mammals will be the most impacted and lose most of their suitable habitats, especially in highlands. The scenario is even worse in the eastern half of Caatinga where habitat destruction already prevails, compounding the threats faced by species there. While species-specific responses can vary with respect to dispersal, behaviour, and energy requirements, our findings indicate that climate change can drive mammal assemblages to biotic homogenisation and species loss, with drastic changes in assemblage trophic structure. For successful long-term socioenvironmental policy and conservation planning, it is critical that findings from biodiversity forecasts are considered.

## INTRODUCTION

Defaunation typically refers to the depletion of fauna caused by overexploitation, habitat destruction, and invasive species (Dirzo et al., 2014; Redford, 1992). At large spatial scales, defaunation may generate complex spatial patterns rather than a simple reduction in species richness, which depend on species-specific responses to defaunation drivers and landscape configuration (Bogoni et al., 2020) Climate change adds another layer of complexity to the spatial consequences of defaunation, since, besides posing an additional threat to wildlife, it is expected to reshape species distribution patterns. In response to a changing the climate, species can be displaced to regions with more favourable conditions, experiencing either geographic range contraction or expansion (Lenoir & Svenning, 2015). Species with higher tolerance to environmental change (e.g., disturbance-adapted, habitat generalists, wide-ranging, and synanthropic species) are less likely to be affected and may even expand their occurrence to novel habitats. In contrast, more sensitive species (e.g., habitat specialists, narrow-ranging species) may lose suitable areas and eventually becoming locally extinct (Filgueiras et al., 2021). These differences in species responses have the potential to change richness and composition of local assemblages, ultimately affecting biodiversity patterns.

The widespread loss of specialist species reduces local species richness (alpha diversity) and may increase the similarity in species composition across space, decreasing beta diversity, a phenomenon termed biotic homogenisation (Mckinney & Lockwood, 1999). Most often, biotic homogenization also result from increases in local richness due to the colonization of species assemblages by generalists (Filgueiras et al., 2021; Socolar et al., 2016). However, species redistribution may also increase the spatial heterogeneity in assemblage composition, either due to the gain of disturbance-adapted species or to the loss of widespread species (Socolar et al., 2016). Although studies on the effects of climate change over biodiversity patterns often emphasize the biotic homogenization due to species loss (Clavel et al., 2011; Hidasi-Neto et al., 2019; Moura et al., 2023), the prevalence each of those process is likely context dependent, and spatial patterns will vary according to species composition, the level of spatial heterogeneity in environmental conditions and the severity of climate changes.

The potential effects of biotic homogenization have been studied mostly in tropical rainforests (Sales et al., 2020), leaving other types of systems highly subject to climate change understudied. Because future climate projections also include changes in the volume, frequency, and geography of rainfalls (IPCC, 2021), climate change is particularly worrying for regions already facing scarcity of water. For instance, drylands are expected to become hotter, drier, and less suitable for a significant portion of their species (Aguirre-Gutiérrez et al., 2020). If these projections are confirmed, it is likely that drylands will gradually become impoverished, homogenised, and driven towards desertification (Moura et al., 2023; Torres et al., 2017). One of Earth’s most vulnerable drylands, the Caatinga, is also the largest tropical dry forest in South America (Banda-R et al., 2016; Hoekstra et al., 2004; Silva et al., 2017). In addition to being affected by chronic disturbances (Antongiovanni et al., 2020), this semiarid region underwent a high degree of defaunation associated with habitat loss and poaching in the past five centuries (Alves et al., 2012; Barboza et al., 2016; Bogoni et al., 2020), showing a high proportion of locally threatened species, including endemic ones (Leal et al., 2005) Besides being an ideal study system to the consequences of climate change on biodiversity distribution patterns, investigating the response of tropical dry forest mammals to climate change can help elucidate impacts of environmental change on dryland biodiversity.

In the Caatinga drylands, about half of the mammal species are non-volant (Carmignotto & Astúa, 2018). Although many of these species are shared with neighbouring biomes (Carmignotto et al., 2012), the composition of Caatinga mammals reflects a complex biogeographic history that has involved periodical expansions and retractions of tropical dry forests across different mountain ranges along the Pleistocene (Silva et al., 2017). On the one hand, Caatinga species have historically experienced high climatic variation (Costa et al., 2018), which may have selected organisms able to keep pace with climate change (Riddell et al., 2021; Schloss et al., 2012). If so, future climate change would have limited influence on species richness and composition of mammal assemblages. However, if Caatinga species are already near their physiological limits (Araújo et al., 2013) or have relied on highland humid enclaves as refuges over evolutionary time (Werneck et al., 2011), further increases in arid conditions could trigger a range shift in these species with consequences for assemblage structure.

Herein, we used ecological niche modelling and community ecology biodiversity metrics to examine potential geographical range shifts of non-volant mammal species in the Caatinga and evaluate impacts of climate change on mammal assemblages. We combined data on species distribution and body mass to investigate projected changes in geographical patterns of mammal richness and spatial dissimilarity across different future climate scenarios. Specifically, we sought to determine whether the balance between potential range contraction or expansion may increase or decrease species richness (alpha diversity) and how those changes in distribution may impact homogenisation or heterogenisation of faunal composition (beta diversity) across space. Because ecological losses are often non-random, with large-sized and longer-lived non-volant mammals disappearing first (Carmona et al., 2021; Cooke et al., 2019), we also examined how changes in average body mass per assemblage (if any) was linked to species loss and biotic homogenisation. Because the elevational gradient around highlands appears to sustain more favourable conditions for non-volant mammals (Becker et al., 2007), we expected relatively lower changes in species richness and composition of mammal assemblages at higher elevations, with overall decline in richness and biotic homogenization associated with a reduction in average body mass.

## METHODS

### Species Data

We compiled occurrence data of Caatinga non-volant mammals searching for different term combinations: “mamíferos”, “caatinga”, “nordeste”, “dataset”, “northeast”, “dryland”, and “mammals” in Google Scholar, identifying 185 mammal species known to occur in the Caatinga. We then used 19 published studies to extracted occurrence records collected between 1957 and 2021 (Asfora et al., 2011; Brennand et al., 2013; Culot et al., 2019; Feijó & Langguth, 2013; Freitas, 1957; Gardner, 2008; Geise et al., 2010; Gurgel-Filho et al., 2015; Lima et al., 2017; Malcher et al., 2017; Mares et al., 1981; Mendonça et al., 2018; Nagy□Reis et al., 2020; Nascimento & Feijó, 2017; Oliveira et al., 2003; Patton et al., 2015; Pires & Wied, 1965; Santos et al., 2019; Souza et al., 2019). We also incorporated data from the mastozoological collection of Universidade Federal da Paraíba (the largest mammal collection of Northeastern Brazil) and other collections included in the Global Biodiversity Information Facility (GBIF, 2023). We included species occurrence records if information was available on coordinates, collection year, and species taxonomy in agreement with specialized literature (Carmignotto & Astúa, 2018; Feijó et al., 2016; Feijó & Langguth, 2013; Gardner, 2008; Gurgel-Filho et al., 2015; Nascimento & Feijó, 2017; Oliveira & Langguth, 2004; Patton et al., 2015; Quintela et al., 2020). After excluding the bat species, our database summed 39,459 occurrence records for 93 species of non-volant mammals.

We used the *CoordinateCleaner* R package (Zizka et al., 2019) to remove duplicates and geoprocessing errors (records distant less than 1 km from municipality, state, or country centroids, or located over water), leading to 18,758 records. To reduce the potential effect of sampling bias and spatial autocorrelation in the occurrence dataset, we randomly filtered one occurrence record for each species within a radius of ∼10 km (Kramer-Schadt et al., 2013). At this point, all species in the database had at least 5 occurrence records. Our final dataset included 11,900 unique occurrence records of 93 species distributed across the Neotropical realm (Fig. S1). Information on mammal body mass was extracted from the *EltonTraits* (Wilman et al., 2014), *Phylacine* (Faurby et al., 2018) and *Combine* databases (Soria et al., 2021) and complemented through specialised literature (see Data Availability for complete sources on body mass data).

### Current and future projections

We used 19 bioclimatic variables from the *WorldClim* v2.1 (Fick & Hijmans, 2017)□ in the spatial resolution of 5 arc-min (∼100 km² pixel) to represent the current climate. The global bioclimatic layers were cropped to the extent of Neotropical realm (i.e., our model’s background). To avoid problems with multicollinearity and reduce the dimensionality of predictor layers, we conducted a principal component analysis on the bioclimatic layers and retained the predictor axes that cumulatively explained 95% of data variation (De Marco & Nóbrega, 2018)□. We projected the linear relationships between raw predictors and principal components onto new layers representing future climate scenarios using the PCA loading coefficients derived from climatic data.

The future climate projections can vary according to different Shared Socioeconomic Pathways (SSPs) that consider distinct paths to greenhouse gas emissions and the human demographic growth (IPCC, 2021). We employed climate projections for the optimistic (SSP 245) and pessimistic (SSP 585) scenarios for the period of 2041-2060 (hereafter 2060) and for the period 2081-2100 (hereafter 2100), both derived from the 6th IPCC Assessment Report (IPCC, 2021). The SSPs were created in agreement with different Generalised Circulation Models (GCMs) that simulate climatic alterations considering various atmospheric processes (IPCC, 2021). To minimise uncertainties about the choice of a particular GCM (Diniz-Filho et al., 2009; Thuiller et al., 2019), we selected the five distinct GCMs, namely: BCC-CSM2-MR, CNRM-CM6-1, IPSL-CM6A-LR, MIROC6, and MRI-ESM2-0.

### Ecological niche models

Recent investigations have showed that 17 occurrence records would be necessary to build traditional ecological niche models (ENMs) for species in the Caatinga (Sampaio & Cavalcante, 2023; van Proosdij et al., 2016). Because almost 20% of mammal species herein considered did not reach this occurrence threshold, we separated our dataset into species with either <20 presences (considered as ‘rare’) or ≥20 presences (considered as ‘common’). We then applied the traditional ENM approach to model habitat suitability of common species and used the Ensemble of Small Models (ESM) approach (Breiner et al., 2015) to model the rare species. Before modelling, we established the calibration (accessible) area of each species as a buffer around its occurrence records, with a width size equal to the maximum nearest neighbour distance among pairs of occurrences (Barve et al., 2012). Within each species calibration area, we computed pseudo-absences using the ratio of 0.5 presence-absence for common species and 0.1 for rare species to avoid very unbalanced models while maximising sampling units (Barbet-Massin et al., 2012; Liu et al., 2019). To increment discriminatory and explanatory capacities of models, we allocated pseudoabsences following the environmentally constrained method, based on the lowest suitable region predicted by a climate envelope (Engler et al., 2004; Lobo & Tognelli, 2011).

Considering that the algorithm choice can affect the habitat suitability estimation (Diniz-Filho et al., 2009; Rangel & Loyola, 2012), we computed an ensemble of projections using four algorithms. For the species modelled using the traditional ENM approach, we used the following algorithms: Generalised Linear Models (using linear and quadratic terms), Generalised Additive Models (using smooth terms with three dimensions), Maximum Entropy (using 10,000 background points and default features based on *MaxNet* package; Phillips et al., 2017), and Random Forests (with the *mtry* parameter automatically tuned by growing 1000 trees through *tuneRF* function in *randomForest* package; Breiman, 2001; Liaw & Wiener, 2002). For the species modelled using the ESM approach, we used the Generalised Linear Models, Generalised Additive Models (using smooth terms with two dimensions), and Gradient Boosting Models (using learning rate of 0.1 and 100 trees), and Neural Networks (with 2 hidden layers, and decay parameters of 0; Breiner et al., 2018). For each method and rare species, we obtained the ESM by averaging the habitat suitability of bivariate models weighted by their respective model Somers’ D □D = 2 × (AUC − 0.5)] (Breiner et al., 2015). The ESMs computed for the four abovementioned methods were then used to build an ensemble of projections for each rare species.

When projecting ENMs to new regions or time periods, it is possible to project habitat suitability for conditions outside the range represented by the training data (Elith et al., 2010). To account for the impact of model extrapolation on each species projection, we computed the Mobility-Oriented Parity (MOP) metric (Owens et al., 2013) within the calibration area of each species. We calculated the MOP metric by measuring the Euclidean distance between environmental conditions of the projected pixel and the nearest 10% training data observations (Montti et al., 2021). The MOP metric was further normalized to 1 and subtracted from 1 to reflect environmental similarity (Owens et al., 2013). We filtered habitat suitability estimates for projected pixels showing very high (MOP values ≥ 0.9), high (MOP ≥ 0.8), and moderate (MOP ≥ 0.7) environmental similarity with the training data. To minimise issues with unlimited dispersal, we restricted all projections to the respective calibration area defined for each species.

We calibrated the models using 5-folds cross-validation, with 80% of randomly selected observations (presences and pseudo-absences) used for training, and the remaining 20% used for testing at each iteration (Roberts et al., 2017). Model performance was evaluated through computation of Sorensen similarity index (ranging from 0 to 1) between observations and binary predictions (Leroy et al., 2018)□. The habitat suitability threshold selected to make predictions binary was chosen to maximise the Sorensen index. We also computed complementary metrics of model performance, True Skill Statistic (TSS, ranging from −1 to 1) and Area Under Curve (AUC, ranging from 0 to 1) (Liu et al., 2011), to facilitate comparisons across literature. For the current climate, and for each combination of GCM, SSP, and year, we computed the ensemble model as the average weighted habitat suitability across algorithms, with the Sorensen index used as weight (Andrade et al., 2020). The ensemble model was then made binary using average weighted binarization threshold, with weights given by the Sorensen’s index of the respective algorithm (Andrade et al., 2020; Thuiller et al., 2019). We used the standard deviation of habitat suitability across the GCMs as a measure of future model uncertainty.

Lastly, we applied spatial constraints *a posteriori* to minimise overprediction issues associated with species binary maps derived from ENMs. We used the occurrence-based threshold method (OBR) to exclude unreachable patches of current suitable habitats for each species (Mendes et al., 2020). This approach assumes that suitable patches are reachable if they either overlap with species presence records (occupied patch) or are within an edge-edge distance threshold of an occupied suitable patch (Mendes et al., 2020). We defined the distance threshold as the maximum nearest neighbour distance among pairs of occurrences of each species. All computations were performed in R 4.2.0 (R Core Team, 2022) using the *ENMTML* package (Andrade et al., 2020) to build the traditional ENMs and the *flexsdm* package (Velazco et al., 2022) to compute the ESMs.

### Assemblage-level biodiversity metrics

We divided the Caatinga using an equal-area projection grid cell of 10 × 10 km. We overlaid our grid cells (i.e., species assemblages) with binary maps to build presence-absence matrices for the current time and each future scenario (2060 SSP245, 2100 SSP245, 2060 SSP585, and 2100 SSP585). To represent the aggregate model uncertainty in future scenarios, we used the average standard deviation of habitat suitability for species in each grid cell (species assemblage). More specifically, we initially averaged the variances (i.e., the squared deviations) for species habitat suitability in each cell, and then square rooted the outcome to get the average standard deviation (AvgSD) for each future year–SSP scenario combination (2060 SSP245, 2060 SSP585, 2100 SSP245, and 2100 SSP585).

Species richness corresponded to the number of species (*S*) present in each grid cell. The spatial beta-diversity was represented by the multisite Simpson dissimilarity index − β_SIM_ (Baselga, 2010), which is recommended for macroecological investigations given its independence of richness differences (Kreft & Jetz, 2010). We computed β_SIM_ between each focal cell and its immediate neighbouring cells. However, the number neighbouring cells is a proxy to area and can therefore affect the β_SIM_ via species-area relationship (Baselga, 2013). To circumvent this issue, we randomly selected four neighbouring cells around each focal cell to compute β_SIM_. We repeated this procedure 100 times and extracted the average β_SIM_ across iterations to obtain the per cell β_SIM_. Computations were performed in R using the *betapart* package (Baselga & Orme, 2012).

For each grid cell, we also computed the geometric mean of log_10_ body mass across its member species (Avg_mass_) as a proxy for the structure of mammal assemblages (Bogoni et al., 2020). We calculated the richness difference between future and current period (ΔS = S_future_ – S_current_) and change in spatial beta-diversity (Δβ_SIM_ = β_SIM.future_ – β_SIM.current_) to identify species assemblages subject to biotic homogenization (Δβ_SIM_ < 0) or heterogenization (Δβ_SIM_ > 0). Similarly, we computed the ratio of average body mass of future to current projections (MassRatio = Avg_mass.future_ / Avg_mass.current_) to quantify relative changes in mammal assemblages. MassRatio < 1 indicated future assemblage with lower average body mass than today, while MassRatio > 1 indicated the opposite.

To assess the influence of potential topographical refuges in shaping assemblage-level biodiversity metrics in Caatinga, we also categorised grid cells between lowlands (i.e., areas <500 m elevation) and highlands (i.e., areas >500 m elevation). The threshold of 500 meters allowed the detection of the five major Caatinga mountain ranges (e.g., Chapada Diamantina, Planalto da Borborema, Chapada do Araripe, Serra da Ibiapaba, and the highest parts of the Serra da Capivara and Serra das Confusões, see Fig. S2). We used Kruskal-Wallis tests to assess whether the medians of (i) Current species richness, (ii) ΔS, (iii) Avg_mass.current_, and (iv) MassRatio differed between assemblages subject to biotic homogenisation (Δβ_SIM_ < 0) or heterogenisation (Δβ_SIM_> 0) or located in lowlands versus highlands. Linear relationships between projected changes in species richness (ΔS), changes in spatial beta-diversity (Δβ_SIM_), relative changes in average body mass (MassRatio), and aggregated model uncertainty (AvgSD) were verified through a modified t-test (Dutilleul, 1993) to spatially correct the degrees of freedom of correlation coefficients. Computations were performed in R using the package *SpatialPack* (Osorio et al., 2014).

## RESULTS

Across all non-volant mammal species in the Caatinga, the ensemble models showed moderate to high predictive performance using either the traditional Ecological Niche Modelling approach (median Sørensen similarity index = 0.68, range = 0.52–0.98; median TSS = 0.52, range = 0.12–0.97; median AUC 0.78, range=0.52–0.99) or the Ensemble of Small Models approach (median Sørensen similarity index = 0.60, range = 0.24–0.89; median TSS = 0.6, range = 0.19–0.98; median AUC 0.85, range=0.43–0.99; Fig. S3). Although quantitative differences emerged between the SSP scenarios (SSP245 and SSP585) and year (2060 and 2100), results were qualitatively similar. Therefore, we focused here on projections for 2060 and SSP245, and based on highly similar environmental conditions (MOP values ≥ 0.9), but see the Supporting Information for results on complementary projections.

About 87% of non-volant mammal species were projected to lose suitable areas by 2060, with substantial reductions of suitable areas (i.e., >50% of geographic range loss) occurring mainly inside the Caatinga (Fig. S4). For at least 12 modelled species (12.8%), suitable habitats within the Caatinga were projected to be completely absent by 2060 under the SSP245 scenario (Fig. 1), with this number reaching 28 species (30%) under the pessimistic scenario (SSP585) by 2100 (Figs S5-S6). Our ensemble models projected that four species would currently show suitable habitats only outside the Caatinga, suggesting potential source-sink dynamics for these species (Fig. 1). However, it is worth noting that four out of the five species without projected suitable habitats (*Dasyprocta azarae*, *Gracilinanus microtarsus*, *Mirmecophaga tridactyla*, and *Priodontes maximus*,) lacked occurrence records in the Caatinga, despite being listed in regional checklists (Carmignotto & Astúa, 2018).

**Figure 1.**
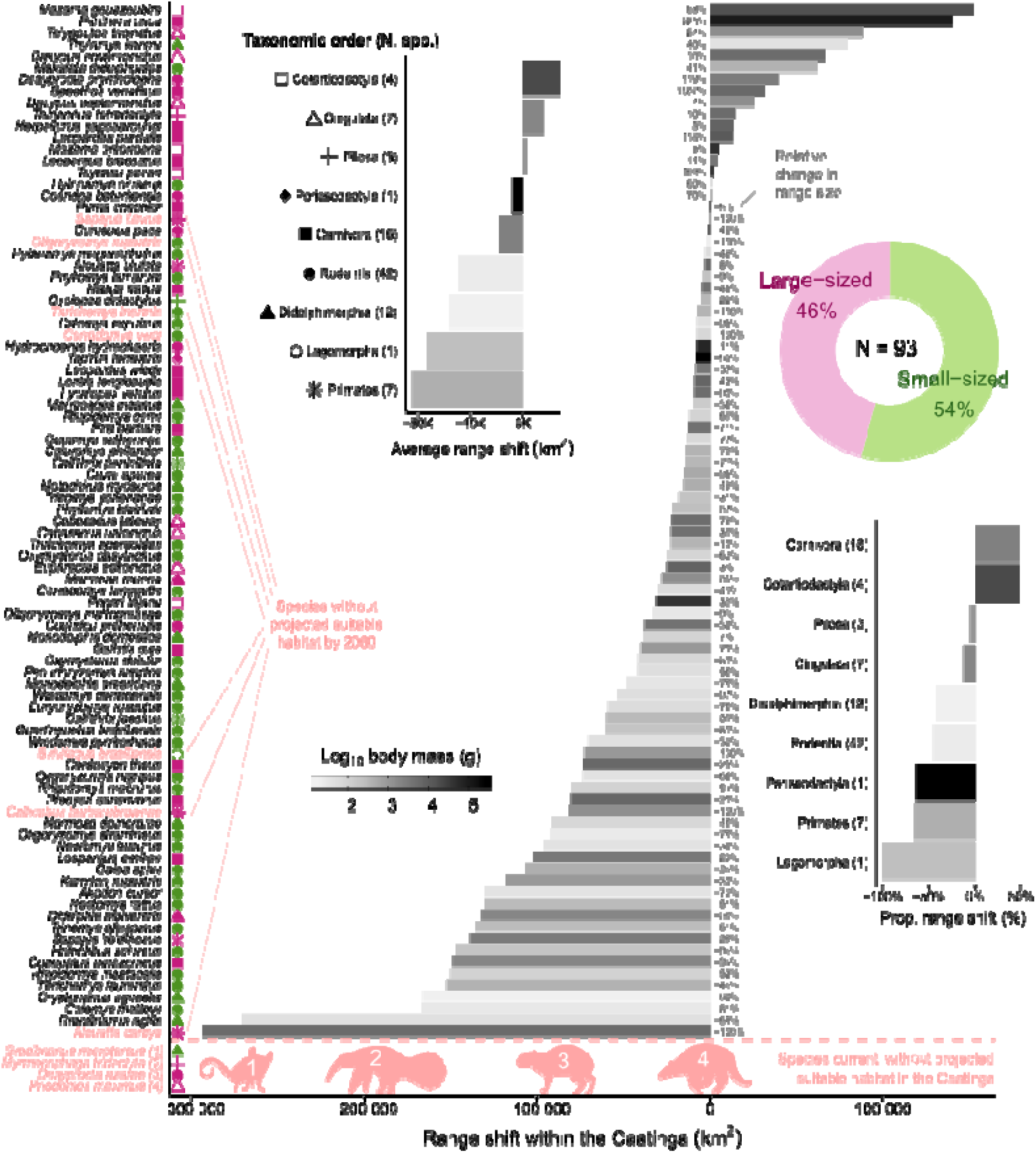
Projected range shift for non-volant mammals in the Caatinga. The four species below the red dashed line showed no current suitable habitats within the Caatinga, although they are projected to occur elsewhere in the Neotropical realm. Species labelled in red elsewhere indicate taxa without projected suitable habitats for 2060 according to the scenario SSP245. Symbol colour on the left panel indicate if species body mass is ≤ 1 kg (green, small-sized) or not (pink, large-sized). Symbol shape follow the taxonomic order indicated in the top-left inset plot. See Figs S4-S6 for results on complementary projections.

Species loss was projected for 91.6% of species assemblages, with an average richness difference of -4.7 species (range ΔS = -23–8) across all assemblages, whereas 69.9% of assemblages showed projected biotic homogenisation (Fig. 2). Median current species richness is higher in regions projected to become more heterogeneous (χ² = 1167.7, d. f. = 7, p < 0.001, Fig. 3a). Similarly, future assemblages projected to be more heterogeneous in the future showed the most pronounced species loss (Fig. S11, Table S1), particularly those in northern Caatinga (Fig. 2), with model uncertainty increasing with richness difference (Fig. S20). Notably, model projections showed low uncertainty across regions subject either to biotic homogenisation or heterogenization (Fig. S21). Assemblages located in lowlands or highlands showed similar changes in species richness and spatial-beta diversity (Figs S12 and S17-18).

**Figure 2.**
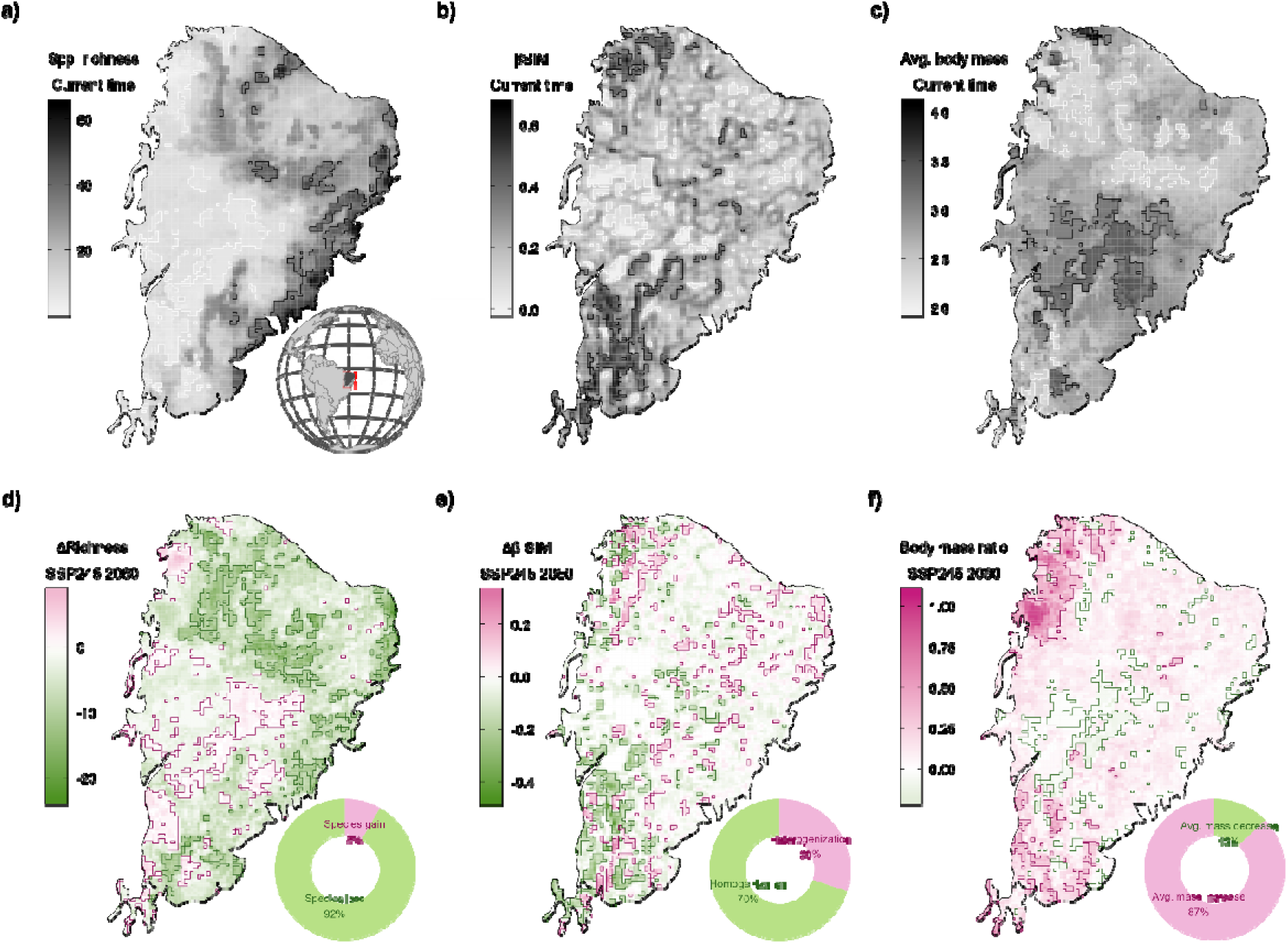
Geographical patterns of species richness, spatial beta-diversity, and average body mass for mammals in the Caatinga. (a) Current species richness, (b) Spatial beta-diversity (β_SIM_), (c) Average log_10_ body mass (g), (d) Projected richness difference (ΔS), (e) Projected change in spatial beta-diversity (Δβ_SIM_), (f) Projected relative change in average body mass. All geographical patterns were derived from species projections holding at least 90% of environmental similarity with training data. The contour lines denote the assemblages (cells) in the upper and lower 10% of the mapped pattern. Plots are shown for the scenario SSP245 at the year 2060. See Figs S7-S10 for results on complementary projections and mapped uncertainty.

**Figure 3.**
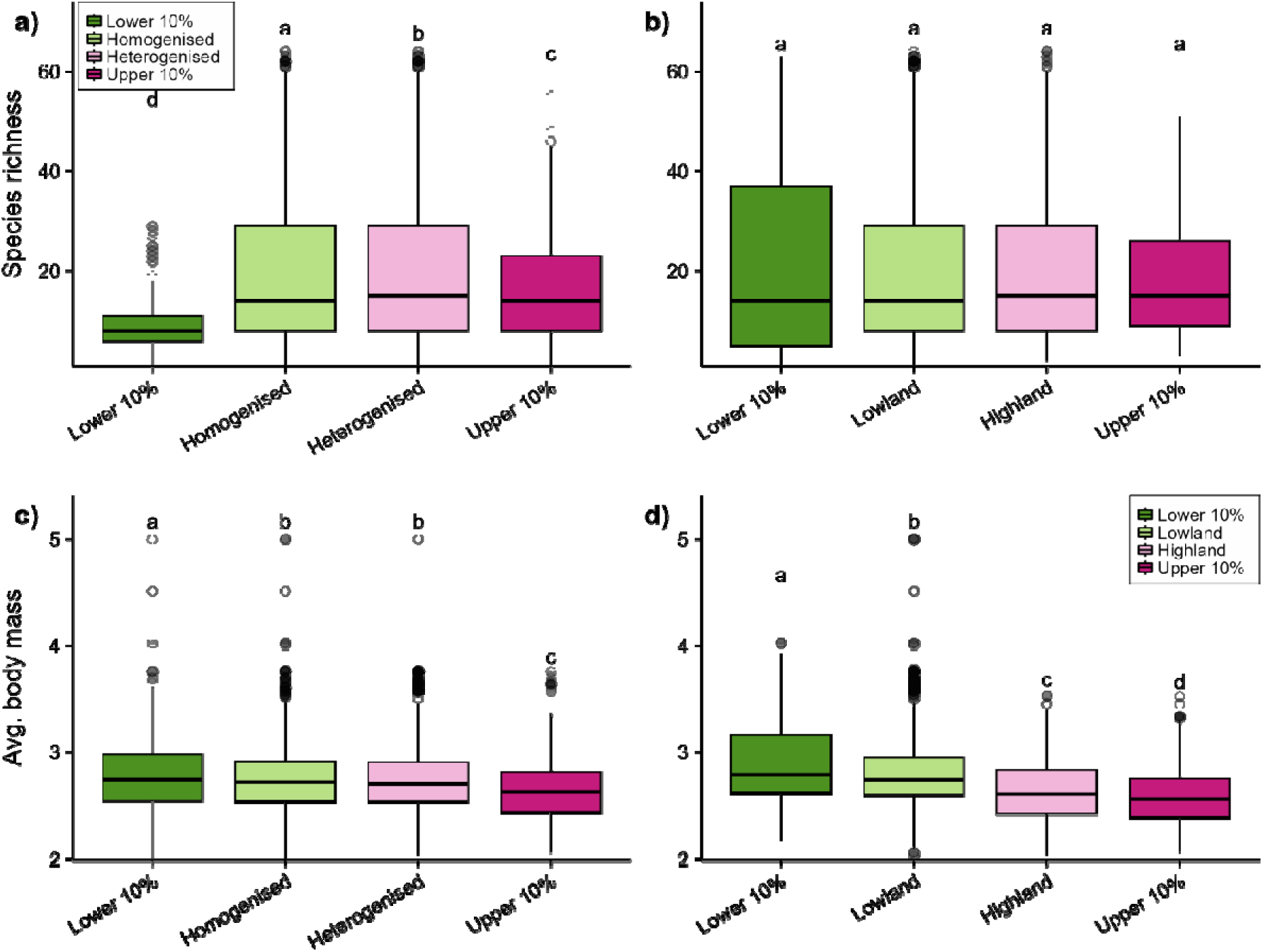
Species richness and average body mass across mammal assemblages at different elevations and levels of biotic change. (a-b) Species richness and (c-d) average body mass. Each box denotes the median (horizontal line), the 25th and 75th percentiles, the 95% confidence intervals (vertical line), and outliers (black dots). Boxplots in darker greenish or pinkish colours denote were computed using the upper and lower 10% assemblages (cells) in terms of biotic change (a, c) and elevation (b, d). Small capital letters denote the results of the Kruskal–Wallis tests for the difference in medians across assemblages subject to different levels of biotic homogenisation or located in lowlands or highlands (boxplots holding the same letter show statistically similar median values under p = 0.05, using Bonferroni correction). Plots are shown for the scenario SSP245 at the year 2060. See Figs S11-S12 and Tables S1-S2 for complementary projections.

Average body mass in current assemblages was generally higher in lowlands than in highlands (χ² = 435.6, d. f. = 7, p < 0.001, Fig. 3c). Surprisingly, 87.7% of assemblages were projected to experience an increase in average body mass of their member species, particularly in the southern and northwestern portions of Caatinga (Fig. 2). The relative change in average body mass was not associated with changes in either species richness (Figs 4c and S14d) or biotic change (Figs 4e and S15d), but tended to slightly increase with elevation (Fig. S12i-l). Across most the SSP scenarios, time periods, and levels of extrapolation constraints, our findings indicated no relationship between changes in average body mass and aggregated model uncertainty (Fig. S22).

**Figure 4.**
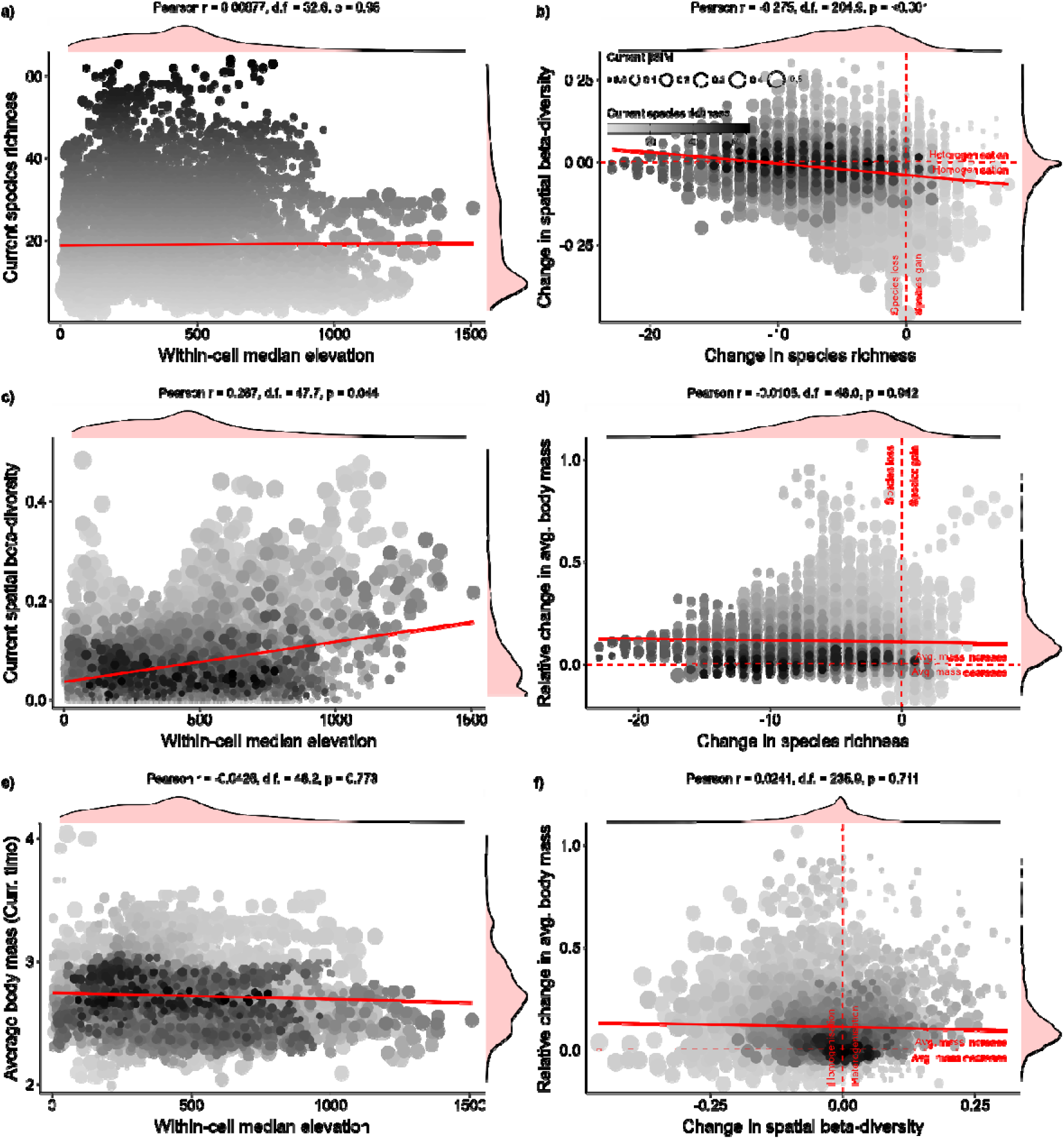
Change in species richness, spatial beta-diversity, average body mass of mammal assemblages in Caatinga. Plots (a, c, e) illustrate the relationship between assemblage-level biodiversity metrics at the current time and elevation, whereas plots (b, d, f) indicate how changes in biodiversity metrics are inter-related. All geographical patterns were derived from species projections holding at least 90% of environmental similarity with training data. Pearson correlations at the top of each panel were based on spatially corrected degrees of freedom. Plots are shown for the scenario SSP245 at the year 2060. See Figs S13-S22 for results on complementary projections.

## DISCUSSION

Drylands in northern South America are expected to face temperature rise of up to 2.7°C by 2060, with changes in the number of consecutive dry days increasing by as much as 21 days (IPCC, 2021). Our study reveals the potential for such changes to drastically erode the diversity of non-volant mammals in the Caatinga. Our projections indicate that most species will lose suitable environmental conditions within the Caatinga, while a few will expand their distribution, which will result in lower species richness and increased compositional similarity to nearby assemblages. Our results show that the biotic homogenisation and species loss are projected in opposite directions, with species gain occurring mostly in regions that are currently species-poor. Although the current beta-diversity is higher in highlands than lowlands, projected changes in biotic composition are only weakly or not at all associated with elevation. Most assemblages are expected to lose small-sized mammals, while large-sized species are projected to colonise neighbouring assemblages. Overall, we reveal how climate change strengthen the defaunation of non-volant mammals and produce complex spatial patterns in the largest tropical dry forest of South America.

Despite mammal adaptations to survive in drylands (e.g., insectivorous diet, night activity, and subterranean shelters), climate change can restrict their physiology and fitness by increasing dehydration, overheating, starvation, and reducing reproduction (Fuller et al., 2021). The projected loss of suitable habitat for almost 90% of all non-volant mammals of Caatinga suggests that these species will have to cope with extreme climate conditions for their dispersion across the biome. Among the main climatic “losers”– species with greatest suitable habitat loss – are primates and the Brazilian cottontail rabbit, but several species from the orders Didelphimorphia and Rodentia also emerge, such as the agile gracile opossum (*Gracilinanus agilis*), the long-tailed climbing mouse (*Rhipidomys mastacalis*), and the white-spined Atlantic spiny-rat (*Trinomys albispinus*). These animals are typically small-sized and occupy lower trophic levels, usually producing a greater number of offspring when compared with large-sized mammals (Carmignotto & Astúa, 2018; Feijó & Langguth, 2013; Santini et al., 2013). The few “winner” species include the brown brocket (*Mazama gouazoubira*), the black-rumped agouti (*Dasyprocta prymnolopha*), which have a wide-ranging distribution and a large body size (Carmignotto & Astúa, 2018; Hetem et al., 2014; Santini et al., 2013), and different species of armadillos, which generally have wide tolerance to warm-dry climates (Soibelzon, 2019). These examples illustrate how animals with low vagility can be disproportionally impacted by climate change, which is supported by our findings on the decrease in the relative contribution of small-sized species across mammal assemblages.

The drastic species loss projected for the assemblages of non-volant mammals can be attributed to changes in dispersal, behaviour, and resource availability due to increasing aridity (Marengo et al., 2017; Torres et al., 2017). Firstly, increased aridity can shorten the optimal period for foraging and breeding (Hetem et al., 2014), and ultimately impact the ecological fitness and maintenance of mammal populations (Fuller et al., 2021). Secondly, geographical barriers may further restrict dispersal and hinder access to suitable habitats (Fuller et al., 2021). Thirdly, hotter and dryer conditions can reduce aboveground biomass (Rito et al., 2017; Souza et al., 2019) and alter floristic composition (Rito et al., 2017; Vieira et al., 2022), thereby impacting competition for food resources not only to herbivores, but also to omnivores and carnivores (Marinho et al., 2020; Oliveira & Diniz-Filho, 2010)□□. Since mammals can exhibit size-dependent variation in vagility, behaviour, and energy needs (Ramesh et al., 2015; Santini et al., 2013; Shipley et al., 1994), prolonged periods of heat and droughts can trigger heterogeneous species responses and enhance negative biotic interactions, ultimately leading to the depletion of faunal assemblages.

The predominance of highly-vagile large-sized species across lowland assemblages and the faster turnover of small-sized species in highlands help to explain the increase in mammal beta-diversity along elevational gradients in the Caatinga□ (Lopez et al., 2016; Melo et al., 2009). While small-sized mammals certainly occur across Caatinga lowlands, the future homogenisation is expected to be primarily driven by the loss of suitable habitats for typically small-sized mammals − adults weighting ≤ 1 kg, *sensu* Chiarello (2000) − which constitute 54% of species in the region (Fig. 1). The current predominance of small-sized mammals across highlands can be related to species persistence through elevational range shift across time (Chen et al., 2011)□, which is especially important in the Caatinga due to its climate instability when compared with other regions in South America (Costa et al., 2018)□. Therefore, the impoverished and compositionally similar mammal assemblages in the lowlands may have resulted from the historic accumulation of local extinctions in the Caatinga, particularly of small-sized species with low vagility (Schloss et al., 2012).

Ecological niches of large-sized species may have been underestimated due to past hunting and overexploitation (Sales et al., 2022), which could further increase in the relative contribution of large-sized species in shaping mammal assemblage. However, our data entries may have missed species entirely if past defaunation resulted in the extinction of large-sized species in the Caatinga. Ungulates like the tapir, peccaries, and different deer species that had wider ranges before European colonization are considered locally extinct across most regions within the biome limits (Barboza et al., 2016). The largest extant mammal in most sites, and the ones projected to increase in range are armadillos, which can be very resilient and often thrive in human-modified landscapes (Bovo et al., 2018; Magalhães et al., 2023), with most small mammals including rodents and marsupials projected to undergo range contractions while the potential range of some of the larger-bodied extant species are projected to increase, the average body mass increases as well. In that sense the pattern we found of increasing mean average body mass is the consequence of the expansion of opportunistic species as well as a legacy of past defaunation. It is worth noting that while the geographical pattern of average body mass indicates a general increase in the relative contribution of large-sized species, intraspecific responses may cause mammal body size to decrease in response to a warming climate (Gardner et al., 2011; Villar & Naya, 2018).

While methodical choices and theoretical limitations like climate uncertainty, dispersal limitations, niche conservatism and model transferability (Barve et al., 2012; Diniz-Filho et al., 2009; Guisan & Thuiller, 2005; Owens et al., 2013; Thuiller et al., 2019) may have affected our projections, wee minimized these issues by offering an ensemble of projections across various modelling algorithms (Araújo & New, 2007). We also implemented an ensemble of future projections across different generalized circulation models and future scenarios of climate change (Diniz-Filho et al., 2009; Thuiller et al., 2019). We also applied species-specific spatial restrictions to remove unreachable patches of projected suitable habitats and minimise overprediction issues related to unlimited dispersal by constraining projections to species-specific calibration areas (Mendes et al., 2020). In addition, assumptions of niche conservatism are likely applicable to mammals in the Caatinga, as the upper limits of mammal thermal tolerance are highly conserved in tropical species (Araújo et al., 2013; Khaliq et al., 2015). To minimise model transferability issues, we constrained habitat suitability estimates to environmental conditions similar to those in the training data (Owens et al., 2013). Although the models used in this study varied quantitatively, the projected changes consistently pointed in the same direction, conveying a unified message.

Our findings indicate a higher species loss for mammal assemblages in the eastern half of Caatinga, which is also affected by chronic disturbances (Antongiovanni et al., 2020). The highly fragmented and diminished vegetation cover of eastern Caatinga (Castanho et al., 2020) impose additional challenges for non-volant mammals to track suitable habitats (Alves et al., 2020), further contributing to depauperate the trophic structure of species assemblages (Mendoza & Araújo, 2019). Although mammal assemblages subject to high species loss exhibit more future uncertainty, a more optimistic outlook is unlikely as these regions also overlap with heavily settled human-modified landscapes in the Caatinga (Antongiovanni et al., 2018, 2020) and regions projected to vegetation complexity and diversity (Moura et al., 2023). Therefore, the severe defaunation of Caatinga mammal assemblages is a probable outcome, with small-sized species loss driven by climate change − at least partially − and the depauperating of large-sized mammal further exacerbated by overexploitation and habitat destruction (Alves et al., 2023; Bogoni et al., 2020). In the long-term, this drastic simplification of mammal assemblages can disrupt biotic interactions and impact ecosystem services in tropical dry forests, by reducing the potential for vegetation regeneration and carbon storage (Bello et al., 2015; Fricke et al., 2022; Gardner et al., 2019). The success of long-term socioenvironmental policy and biodiversity conservation planning necessitates that findings derived from biodiversity forecasts are considered.

## ACKNOWLEDGEMENTS

We are grateful to Pedro C. Estrela, Anderson Feijó, Thais Kubik, Daniel P. Silva and Cibele R. Bonvicino for comments on previous versions of this manuscript. To the Mammal Collection from Universidade Federal da Paraíba for providing occurrences data. To Fundação de Amparo à Pesquisa do Estado de São Paulo for grants to MRM (FAPESP #2021/11840-6 and #2022/12231-6) and MMP (FAPESP #2019/25478-7). To Conselho Nacional de Desenvolvimento Científico e Tecnológico for grants (CNPq #312178/2019-0 and #307260/2022-4) and to Universidade Federal da Paraíba (PVA-13357-2020) for grants to BAS. To Re:wild and Dimensions Sciences Bridges for grants to GAO. To Fundação de Amparo à Pesquisa do Estado de Minas Gerais for grants to APP. To Coordenação de Aperfeiçoamento de Pessoal de Nível Superior for a scholarship to GAO and fellowship to APP.

## AUTHOR CONTRIBUTIONS

MRM, GAO, and BAS conceived the study; GAO and APP compiled the data; MRM analysed the data. MRM developed the figures and led the writing. All authors contributed critically to the drafts and gave final approval for publication.

## SUPPLEMENTARY MATERIAL

Supplementary Material is available for this manuscript, including Supplementary Tables (S1–S2) and Supplementary Figures (S1–S22).

