## Supplementary Information for "Climate change should drive mammal defaunation in tropical dry forests"

### Climate change-driven biotic homogenisation of non-volant mammals in tropical dry forests

**This PDF file includes:**

Supplementary Tables

Supplementary Figures

#### SUPPLEMENTARY TABLES

**Table S1. Kruskal–Wallis tests for the difference in medians across assemblages subject to different levels of biotic change.** The results are shown for 2060 and 2100 under four levels of extrapolation constraints. MOP00, MOP70, MOP80, and MOP90 – respectively indicate species-level projections including regions with at least 00%, 70%, 80%, and 90% of environmental similarity with the training data. All p-values had their significance corrected using Bonferroni correction.

| Year | Variable | $\chi^2$ | d.f. | p-value | Extrapolation constraints |
|---|---|---|---|---|---|
| 2060 | Current species richness | 1113.11 | 7 | 0.0000 | MOP00 |
| 2060 | Delta richness (ΔS) | 815.67 | 7 | 0.0000 | MOP00 |
| 2060 | Average body mass (current time) | 197.37 | 7 | 0.0000 | MOP00 |
| 2060 | Body mass ratio | 805.70 | 7 | 0.0000 | MOP00 |
| 2100 | Current species richness | 1273.31 | 7 | 0.0000 | MOP00 |
| 2100 | Delta richness (ΔS) | 1328.58 | 7 | 0.0000 | MOP00 |
| 2100 | Average body mass (current time) | 294.18 | 7 | 0.0000 | MOP00 |
| 2100 | Body mass ratio | 2180.26 | 7 | 0.0000 | MOP00 |
| 2060 | Current species richness | 1165.94 | 7 | 0.0000 | MOP70 |
| 2060 | Delta richness (ΔS) | 752.19 | 7 | 0.0000 | MOP70 |
| 2060 | Average body mass (current time) | 237.54 | 7 | 0.0000 | MOP70 |
| 2060 | Body mass ratio | 765.08 | 7 | 0.0000 | MOP70 |
| 2100 | Current species richness | 1339.03 | 7 | 0.0000 | MOP70 |
| 2100 | Delta richness (ΔS) | 1277.08 | 7 | 0.0000 | MOP70 |
| 2100 | Average body mass (current time) | 307.86 | 7 | 0.0000 | MOP70 |
| 2100 | Body mass ratio | 2127.08 | 7 | 0.0000 | MOP70 |
| 2060 | Current species richness | 1265.17 | 7 | 0.0000 | MOP80 |
| 2060 | Delta richness (ΔS) | 947.39 | 7 | 0.0000 | MOP80 |
| 2060 | Average body mass (current time) | 325.74 | 7 | 0.0000 | MOP80 |
| 2060 | Body mass ratio | 869.84 | 7 | 0.0000 | MOP80 |
| 2100 | Current species richness | 1336.23 | 7 | 0.0000 | MOP80 |
| 2100 | Delta richness (ΔS) | 1400.55 | 7 | 0.0000 | MOP80 |
| 2100 | Average body mass (current time) | 301.67 | 7 | 0.0000 | MOP80 |
| 2100 | Body mass ratio | 2605.89 | 7 | 0.0000 | MOP80 |
| 2060 | Current species richness | 1167.77 | 7 | 0.0000 | MOP90 |
| 2060 | Delta richness (ΔS) | 733.99 | 7 | 0.0000 | MOP90 |
| 2060 | Average body mass (current time) | 237.00 | 7 | 0.0000 | MOP90 |
| 2060 | Body mass ratio | 765.95 | 7 | 0.0000 | MOP90 |
| 2100 | Current species richness | 1203.01 | 7 | 0.0000 | MOP90 |
| 2100 | Delta richness (ΔS) | 1136.49 | 7 | 0.0000 | MOP90 |
| 2100 | Average body mass (current time) | 263.46 | 7 | 0.0000 | MOP90 |
| 2100 | Body mass ratio | 1869.48 | 7 | 0.0000 | MOP90 |

**Table S2. Kruskal–Wallis tests for the difference in medians across assemblages at different elevations.** The results are shown for 2060 and 2100 under four levels of extrapolation constraints. MOP00, MOP70, MOP80, and MOP90 – respectively indicate species-level projections including regions with at least 00%, 70%, 80%, and 90% of environmental similarity with the training data. All p-values had their significance corrected using Bonferroni correction.

| Year | Variable | $\chi^2$ | d.f. | p-value | Extrapolation constraints |
|---|---|---|---|---|---|
| 2060 | Current species richness | 8.41 | 7 | 0.2982 | MOP00 |
| 2060 | Delta richness (ΔS) | 510.57 | 7 | 0.0000 | MOP00 |
| 2060 | Average body mass (current time) | 2316.66 | 7 | 0.0000 | MOP00 |
| 2060 | Body mass ratio | 369.86 | 7 | 0.0000 | MOP00 |
| 2100 | Current species richness | 8.41 | 7 | 0.2982 | MOP00 |
| 2100 | Delta richness (ΔS) | 222.28 | 7 | 0.0000 | MOP00 |
| 2100 | Average body mass (current time) | 2316.66 | 7 | 0.0000 | MOP00 |
| 2100 | Body mass ratio | 2196.25 | 7 | 0.0000 | MOP00 |
| 2060 | Current species richness | 2.21 | 7 | 0.9470 | MOP70 |
| 2060 | Delta richness (ΔS) | 496.36 | 7 | 0.0000 | MOP70 |
| 2060 | Average body mass (current time) | 1809.65 | 7 | 0.0000 | MOP70 |
| 2060 | Body mass ratio | 428.40 | 7 | 0.0000 | MOP70 |
| 2100 | Current species richness | 2.21 | 7 | 0.9470 | MOP70 |
| 2100 | Delta richness (ΔS) | 176.61 | 7 | 0.0000 | MOP70 |
| 2100 | Average body mass (current time) | 1809.65 | 7 | 0.0000 | MOP70 |
| 2100 | Body mass ratio | 2157.96 | 7 | 0.0000 | MOP70 |
| 2060 | Current species richness | 2.21 | 7 | 0.9470 | MOP80 |
| 2060 | Delta richness (ΔS) | 321.88 | 7 | 0.0000 | MOP80 |
| 2060 | Average body mass (current time) | 1809.65 | 7 | 0.0000 | MOP80 |
| 2060 | Body mass ratio | 783.19 | 7 | 0.0000 | MOP80 |
| 2100 | Current species richness | 2.21 | 7 | 0.9470 | MOP80 |
| 2100 | Delta richness (ΔS) | 126.22 | 7 | 0.0000 | MOP80 |
| 2100 | Average body mass (current time) | 1809.65 | 7 | 0.0000 | MOP80 |
| 2100 | Body mass ratio | 3126.53 | 7 | 0.0000 | MOP80 |
| 2060 | Current species richness | 2.68 | 7 | 0.9127 | MOP90 |
| 2060 | Delta richness (ΔS) | 502.52 | 7 | 0.0000 | MOP90 |
| 2060 | Average body mass (current time) | 1821.51 | 7 | 0.0000 | MOP90 |
| 2060 | Body mass ratio | 435.60 | 7 | 0.0000 | MOP90 |
| 2100 | Current species richness | 2.68 | 7 | 0.9127 | MOP90 |
| 2100 | Delta richness (ΔS) | 200.42 | 7 | 0.0000 | MOP90 |
| 2100 | Average body mass (current time) | 1821.51 | 7 | 0.0000 | MOP90 |
| 2100 | Body mass ratio | 1977.17 | 7 | 0.0000 | MOP90 |

### SUPPLEMENTARY FIGURES

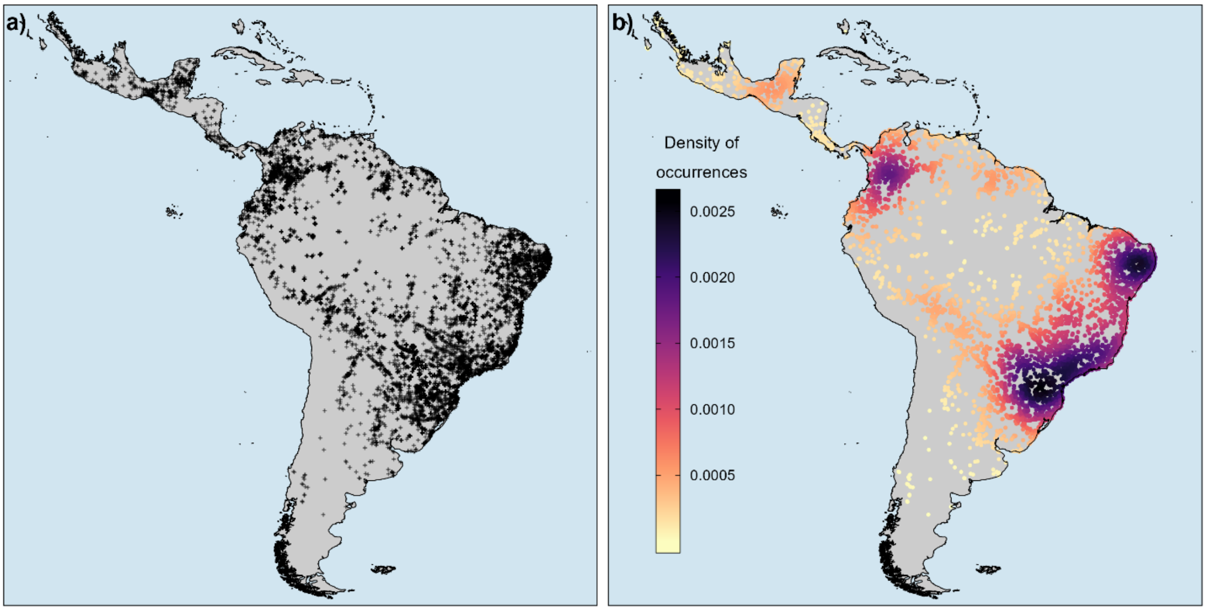


**Fig. S1. Species occurrence records for mammal species known to occur within the Caatinga.** (a) The map shows 11,900 unique occurrence records of 93 species of non-volant mammals. (b) Density map of species occurrence records. The Neotropical realm was the background area used in the modelling procedures.

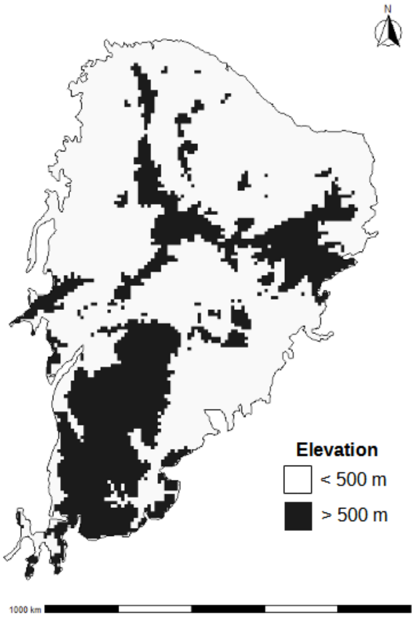


**Fig. S2. Schematic representation of lowland and highland regions in Caatinga.** The threshold of 500 m was used to categorise the 10× 10 km grid cells into high or low elevation.

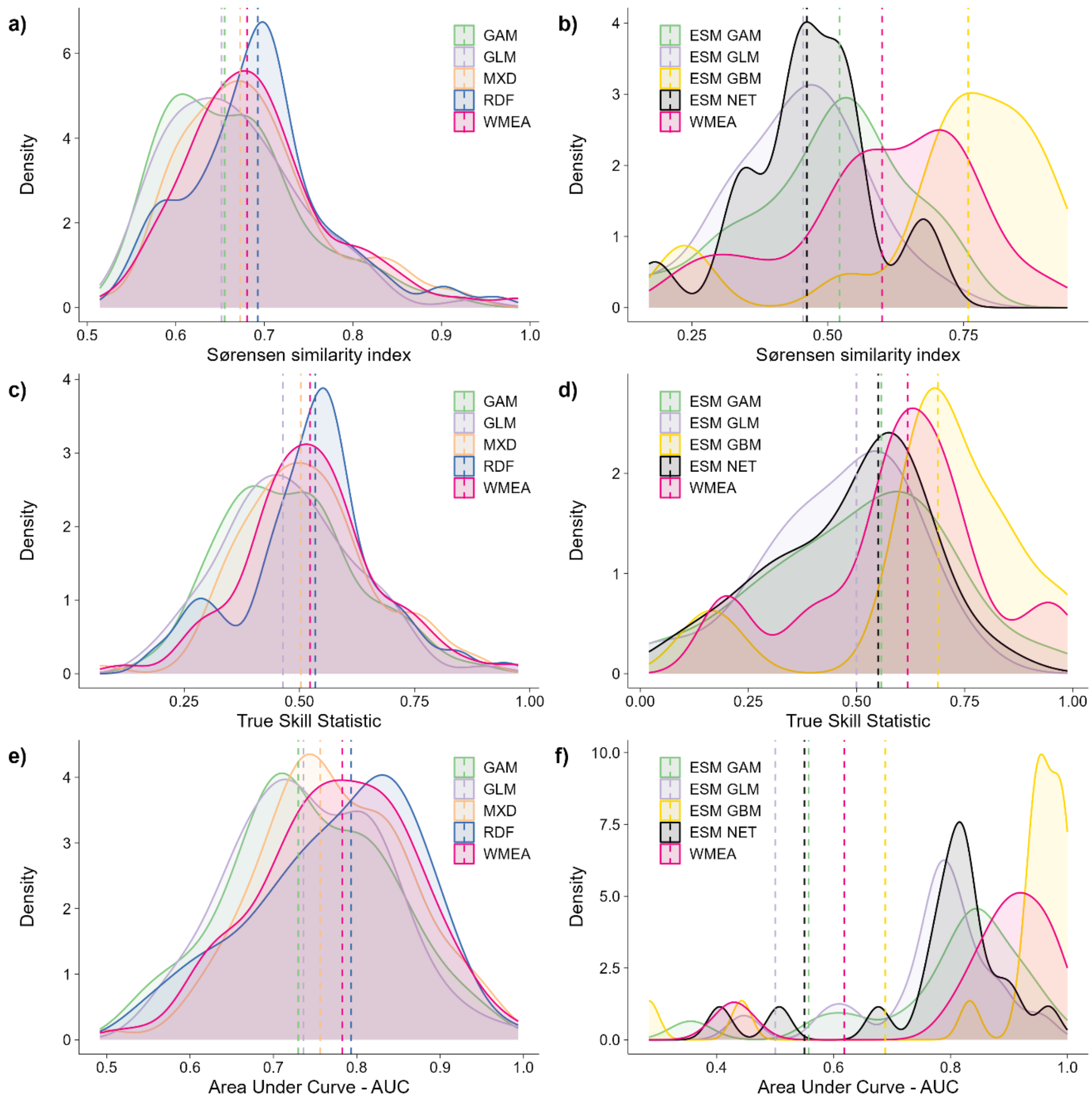


**Fig. S3. Performance metric of different algorithms used to model species distributions.** Plots shows multivariate models built using different algorithms using either the traditional ENM approach (a, c, e) or the ESM approach (b, d, f). Vertical line denotes the mean value of the performance metric observed for the respective algorithm. Algorithm abbreviations: BIO = Bioclim climate envelope, GAM = Generalised Additive Models, GLM = Generalised Linear Models, GBM = Generalised Boosting Regression, MXD = Maximum Entropy, RDF = Random Forests, NET = Neural Network. For each model approach, WMEA represents the ensemble model computed as the weighted average across all available algorithms using the Sørensen similarity index as weight.

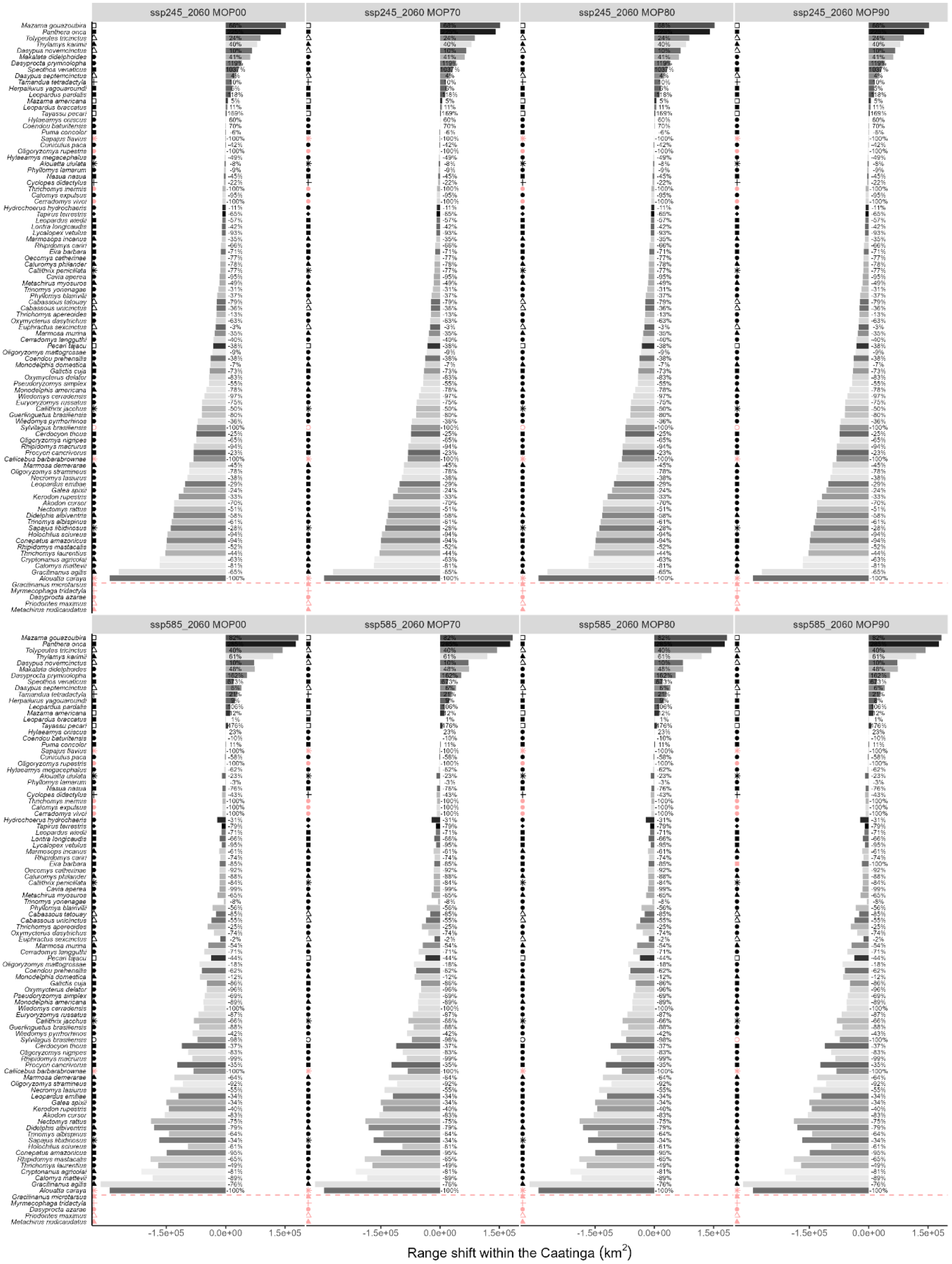


**Fig. S4. Projected range shift for non-volant mammal in the Caatinga for 2060.** Species below the red dashed line showed projected presence only outside the Caatinga, even for the current period. Symbols colours on the left panel indicate species without projected and reachable suitable habitats in Caatinga, whereas symbol shape refers to taxonomic order (see main text for details on each order).

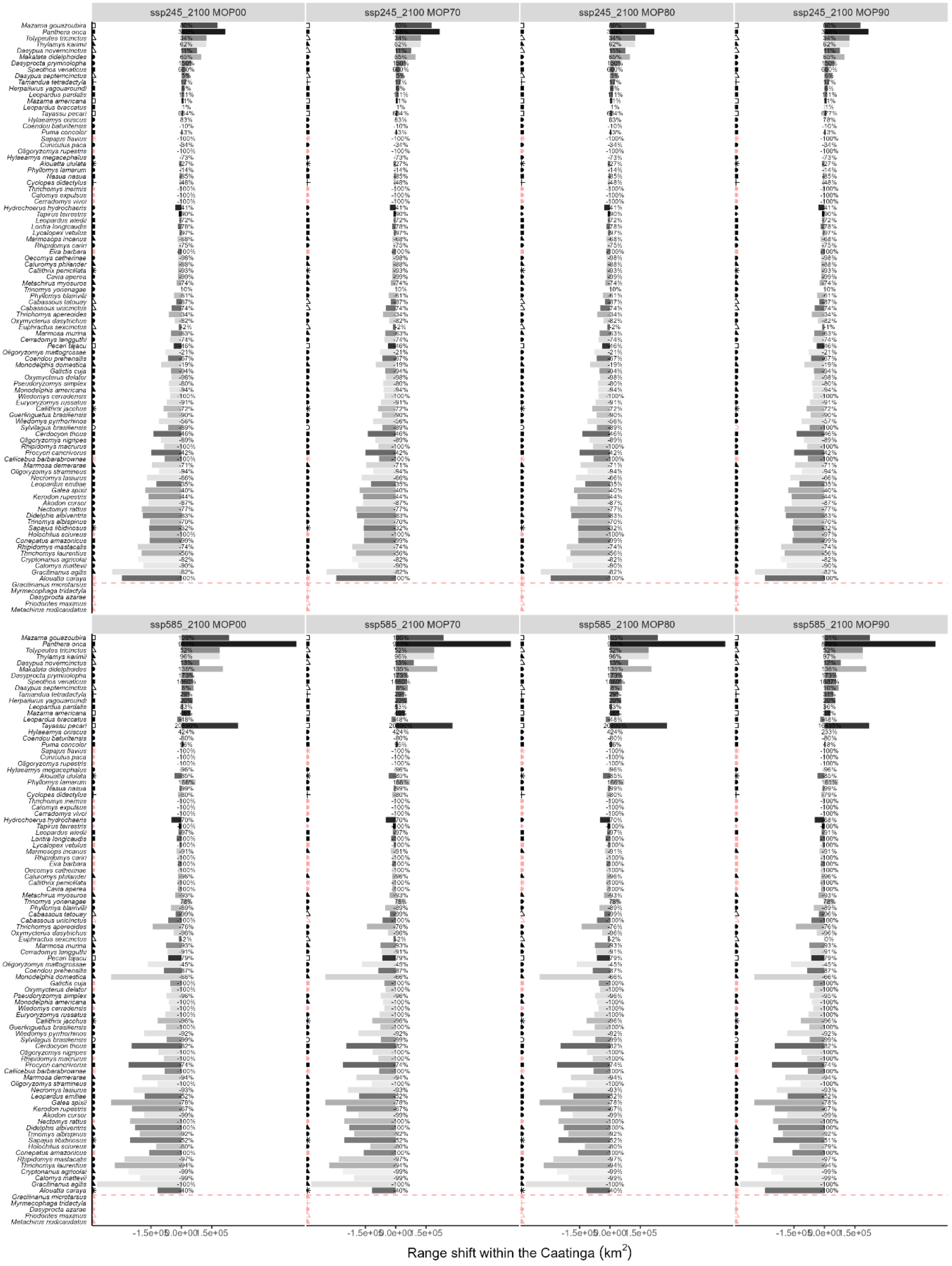


**Fig. S5. Projected range shift for non-volant mammal in the Caatinga for 2100.** Species below the red dashed line showed projected presence only outside the Caatinga, even for the current period. Symbols colours on the left panel indicate species without projected and reachable suitable habitats in Caatinga, whereas symbol shape refers to taxonomic order (see main text for details on each order).

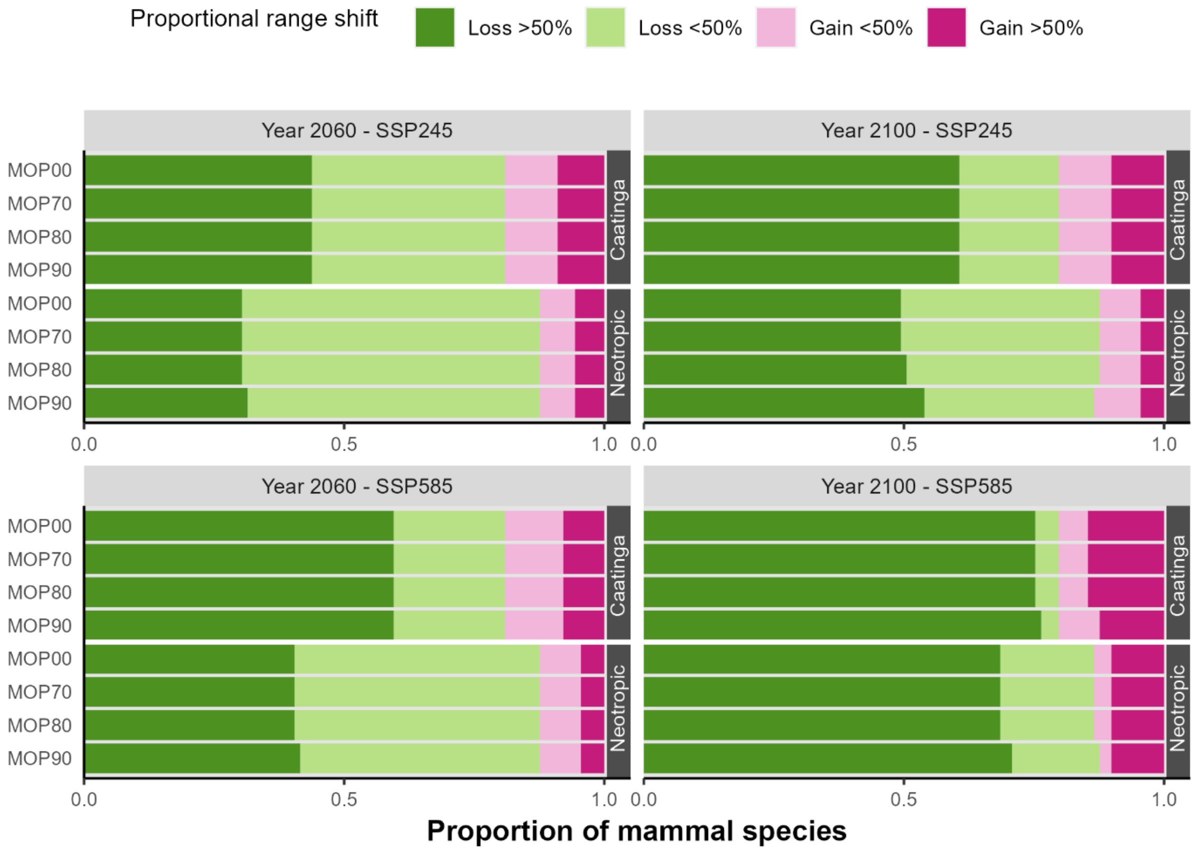


**Fig. S6. Projected range shift for mammal species.** Range shifts were computed separately within the Neotropics and Caatinga. The results are shown for 2060 and 2010 under the business-as-usual (SSP245) and non-mitigation (SSP585) scenarios. The four levels of extrapolation constraints – MOP00, MOP70, MOP80, and MOP90 – respectively indicate species-level projections including regions with at least 00%, 70%, 80%, and 90% of environmental similarity with the training data.

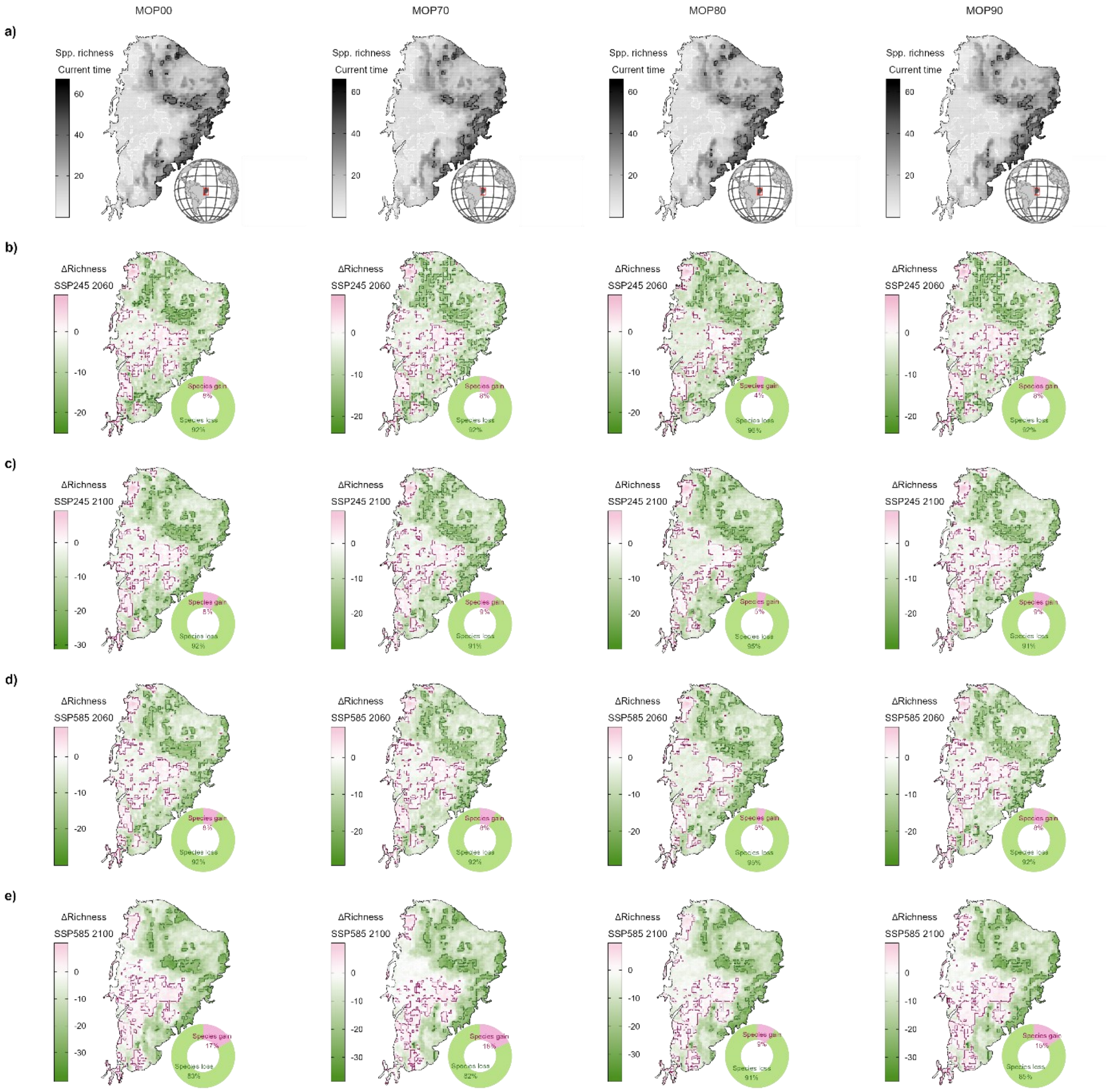


**Fig. S7. Geographical patterns of non-volant mammal richness in the Caatinga.** Current species richness (a) and projected changes in richness (b-e; ΔS) are show for all combinations of future year and SSP scenario. The four levels of extrapolation constraints – MOP00, MOP70, MOP80, and MOP90 – respectively indicate species-level projections including regions with at least 00%, 70%, 80%, and 90% of environmental similarity with the training data. The contour lines denote the assemblages (cells) in the upper and lower 10% of the mapped pattern.

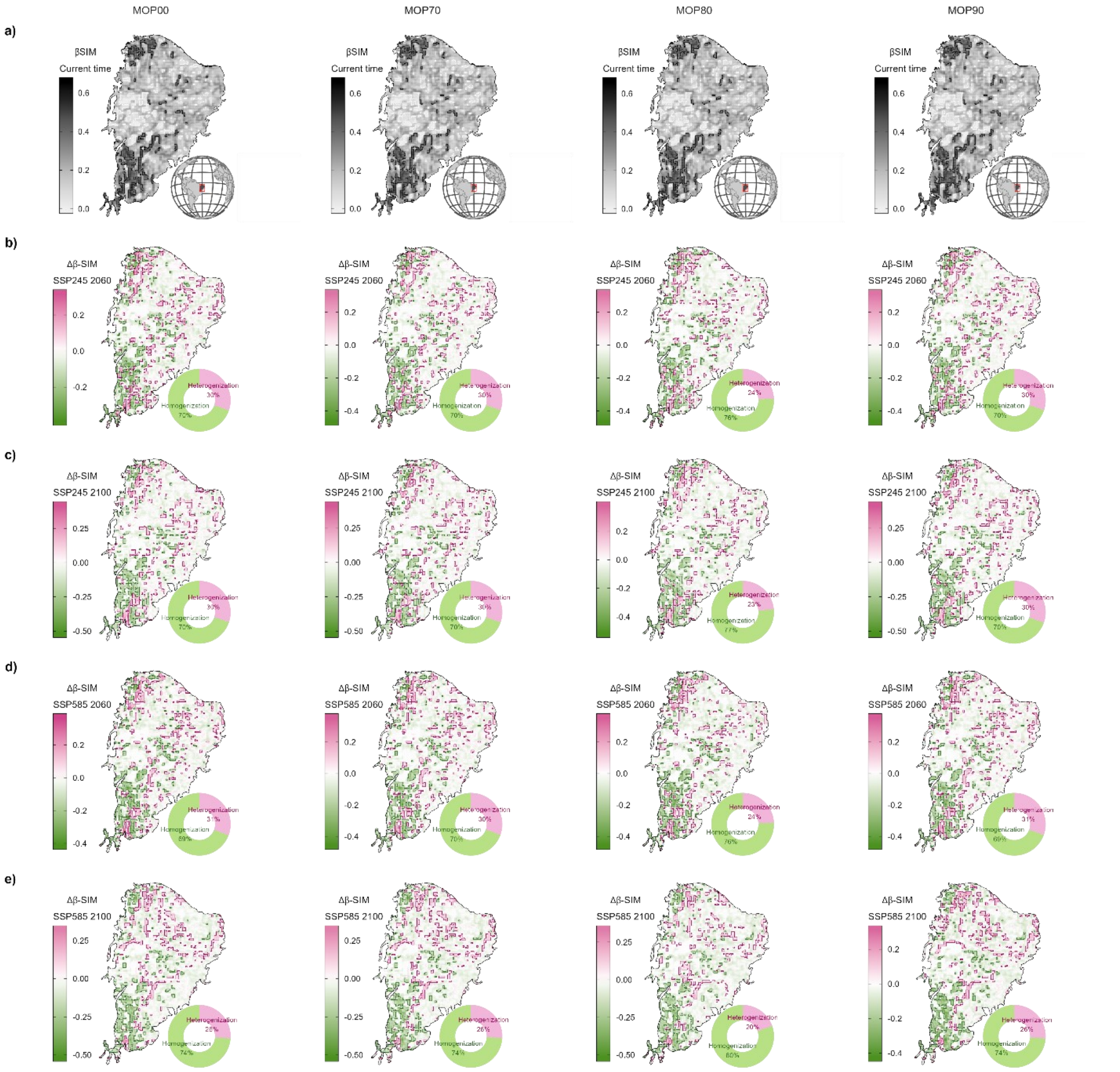


**Fig. S8. Geographical patterns of non-volant mammal beta diversity in the Caatinga.** Current beta-diversity (a) and expected changes in spatial beta-diversity (b-e; $\Delta\beta_{SIM}$) are show for all combinations of future year and SSP scenario. The four levels of extrapolation constraints – MOP00, MOP70, MOP80, and MOP90 – respectively indicate species-level projections including regions with at least 00%, 70%, 80%, and 90% of environmental similarity with the training data. The contour lines denote the assemblages (cells) in the upper and lower 10% of the mapped pattern.

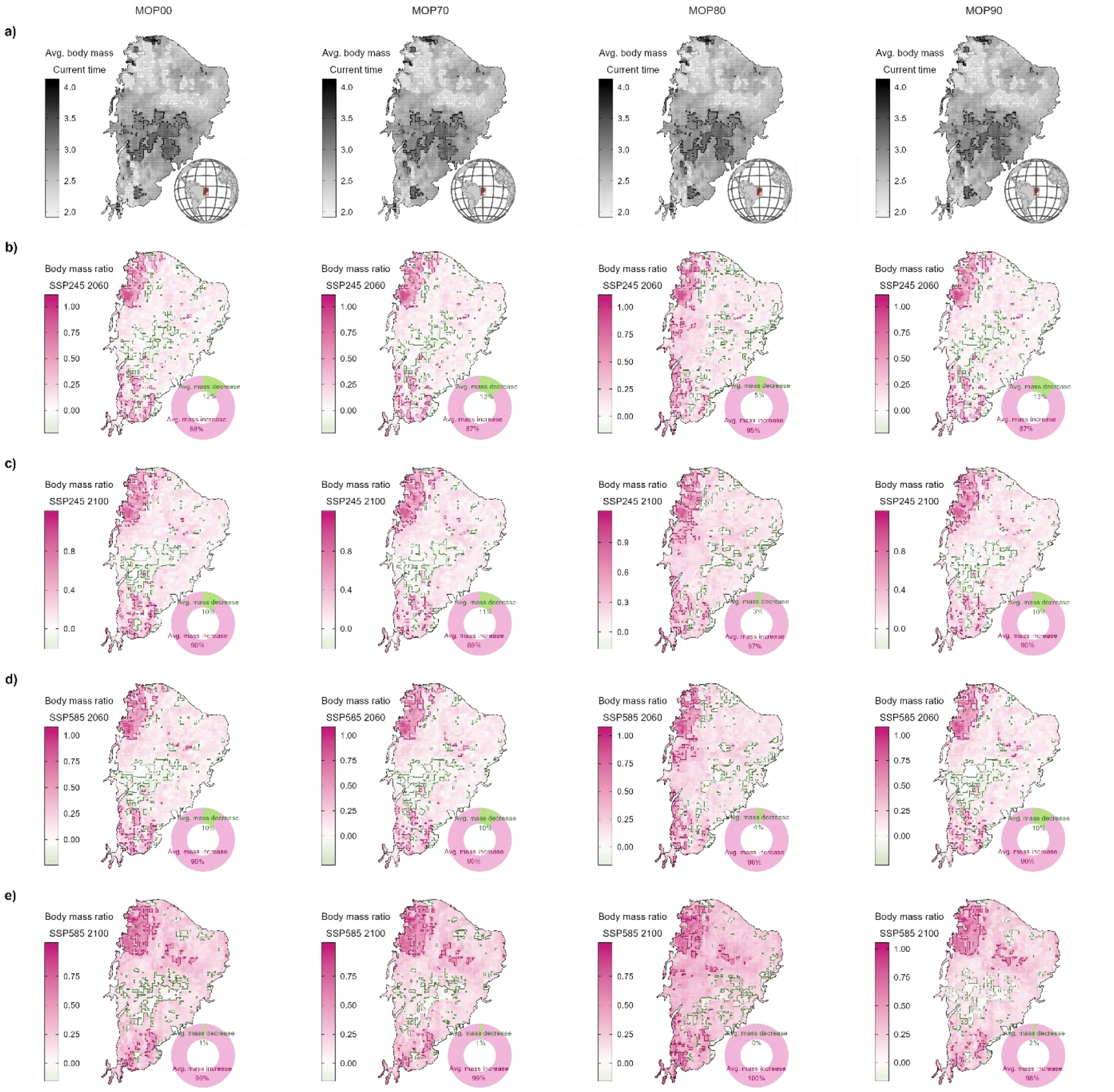


**Fig. S9. Geographical patterns of non-volant mammal body mass in the Caatinga.** Current average body mass (a) and relative change in average body mass (b-e) are show for all combinations of future year and SSP scenario. The four levels of extrapolation constraints – MOP00, MOP70, MOP80, and MOP90 – respectively indicate species-level projections including regions with at least 00%, 70%, 80%, and 90% of environmental similarity with the training data. Maps shows the geometric mean of $\log_{10}$(body mass) of species in each 10 × 10 km cell. We subtracted '1' from the mass ratio metric to centre it around 0. The contour lines denote the assemblages (cells) in the upper and lower 10% of the mapped pattern.

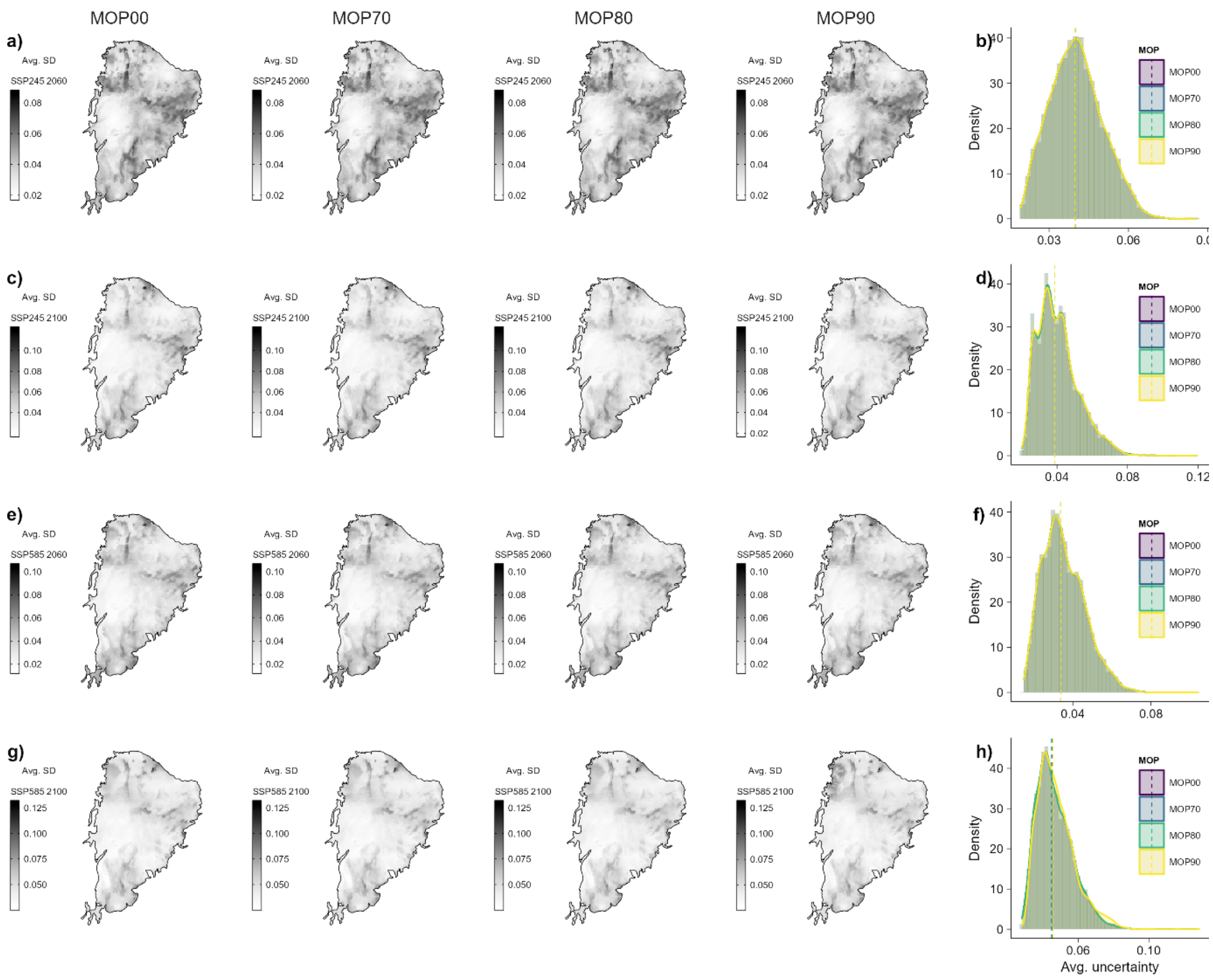


**Fig. S10. Uncertainty of species modelled distributions across generalised circulation models (GMCs) in the Caatinga.** Each map shows the average standard deviation of species habitat suitability across future GCMs. Panel rows (a, c, e, g) provide uncertainty maps for a same year and SSP scenario. The four levels of extrapolation constraints – MOP00, MOP70, MOP80, and MOP90 – respectively indicate species-level projections including regions with at least 00%, 70%, 80%, and 90% of environmental similarity with the training data. Histograms (b, d, f, h) show the frequency of uncertainty values under the four levels of extrapolations allowed.

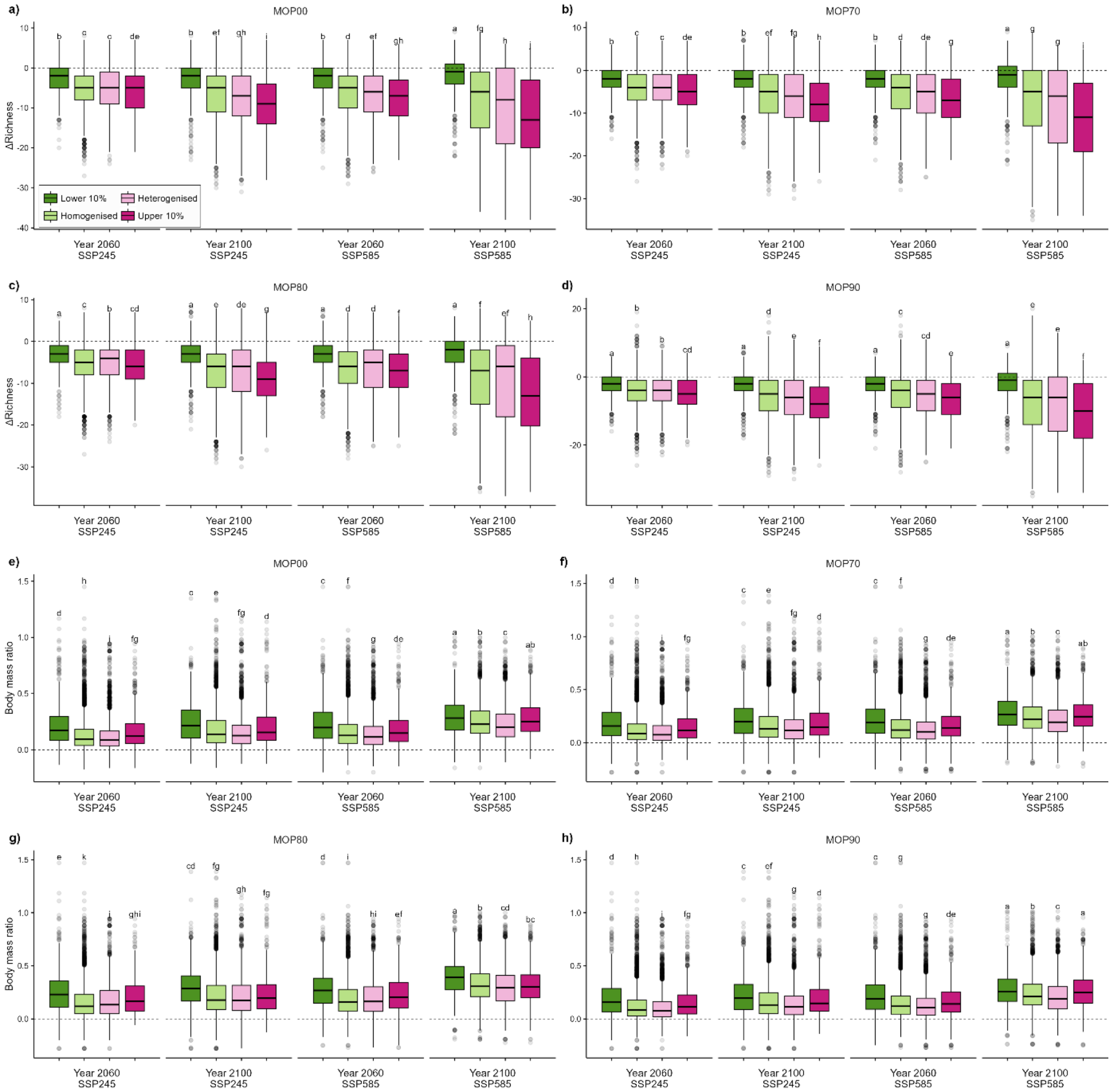


**Fig. S11. Change in assemblage-level metrics across levels of biotic change**. (a-d) Richness difference (ΔS), (e-h) Relative change in average body mass (Mass ratio − 1). Each box denotes the median (horizontal line), the 25th and 75th percentiles, the 95% confidence intervals (vertical line), and outliers (dots). Small capital letters show the results of the Bonferroni corrected Kruskal–Wallis tests for the difference in median assemblage values across levels of biotic change (see Table S1). The four levels of extrapolation constraints – MOP00, MOP70, MOP80, and MOP90 – respectively indicate species-level projections including regions with at least 00%, 70%, 80%, and 90% of environmental similarity with the training data.

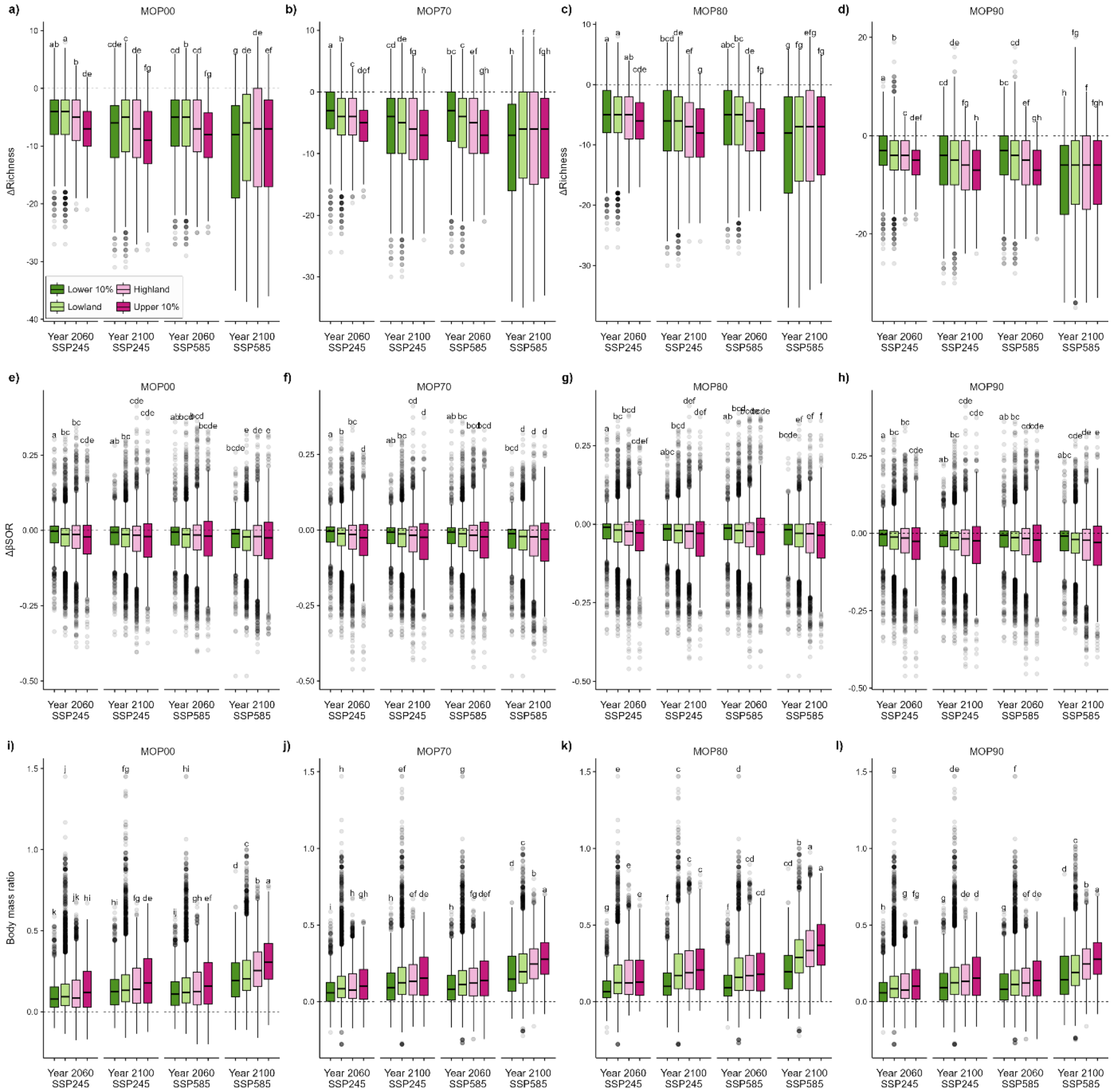


**Fig. S12. Change in assemblage-level metrics between lowlands and highlands**. (a-d) Richness difference (ΔS), (e-h) spatial beta-diversity ($\Delta\beta_{SIM}$), (i-l) relative change in average body mass (Mass ratio − 1). Each box denotes the median (horizontal line), the 25th and 75th percentiles, the 95% confidence intervals (vertical line), and outliers (dots). Small capital letters show the results of the Bonferroni corrected Kruskal–Wallis tests for the difference in median assemblage values across levels of biotic change (see Table S1). The four levels of extrapolation constraints – MOP00, MOP70, MOP80, and MOP90 – respectively indicate species-level projections including regions with at least 00%, 70%, 80%, and 90% of environmental similarity with the training data.

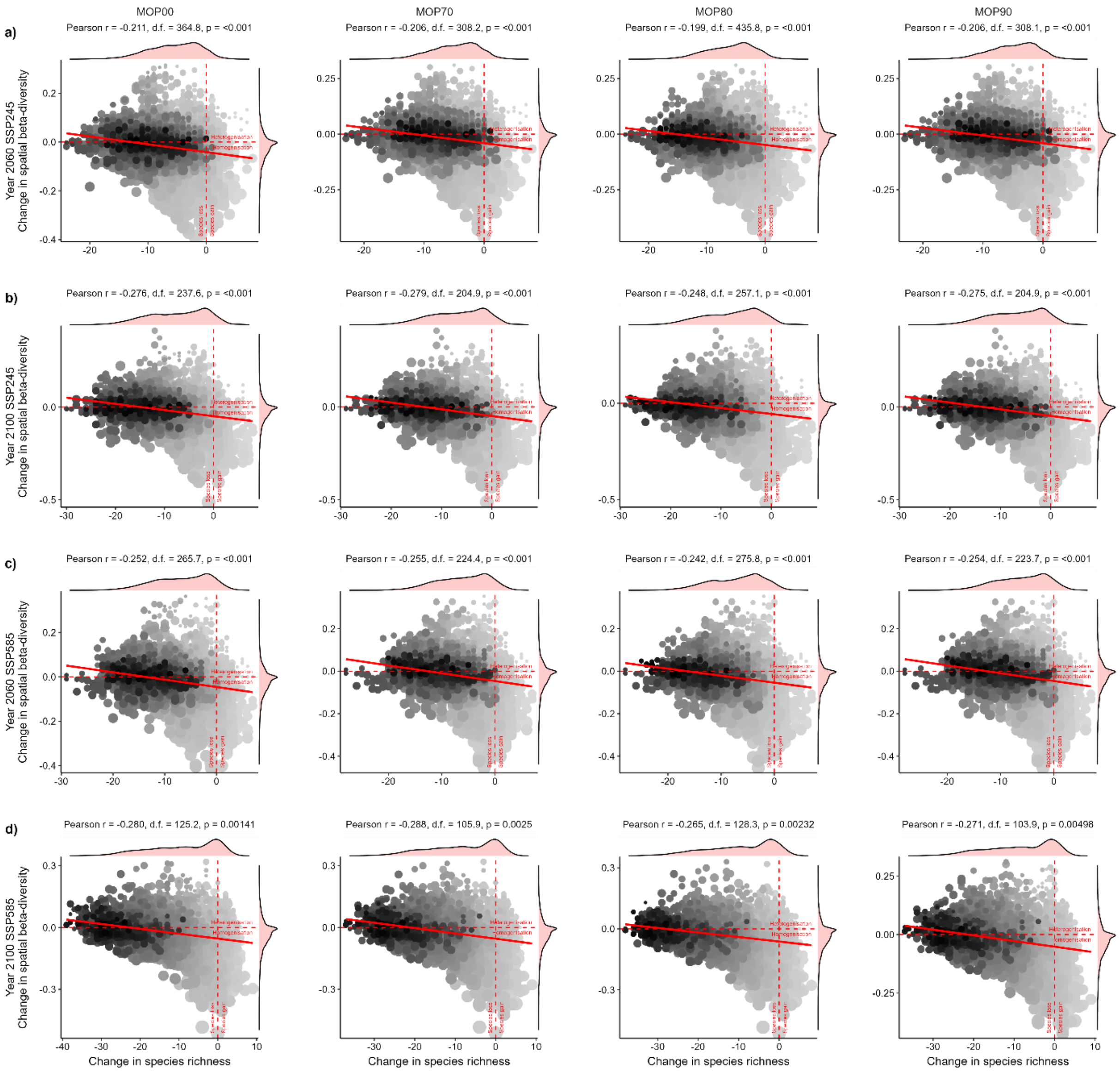


**Fig. S13. Relationship between change in species richness and spatial beta-diversity in Caatinga non-volant mammal assemblages.** Relationship between differences in species richness (ΔS) and spatial beta-diversity ($\Delta\beta_{SIM}$) in the future scenarios (a) SSP245 – 2060, (b) SSP245 – 2100, (c) SSP585 – 2060, (d) SSP585 – 2100. The four levels of extrapolation constraints – MOP00, MOP70, MOP80, and MOP90 – respectively indicate species-level projections including regions with at least 00%, 70%, 80%, and 90% of environmental similarity with the training data. Symbol colours indicate the current species richness (darker = higher richness) per 10 × 10 km cell. Symbol size is proportional to the current spatial beta-diversity ($\beta_{SIM}$). Pearson correlations at the top of each panel were based on spatially corrected degrees of freedom.

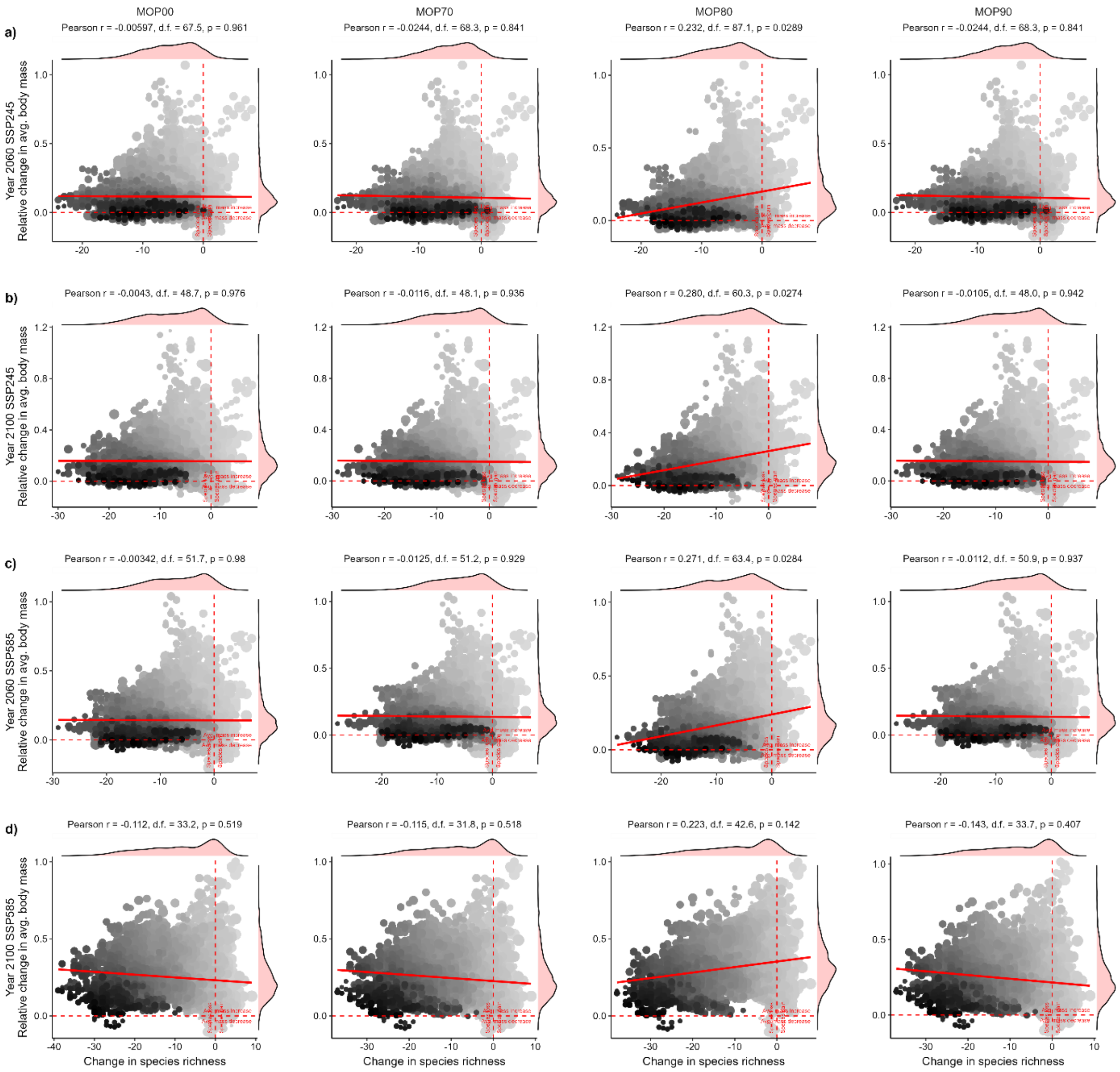


**Fig. S14. Relationship between change in species richness and average body mass in Caatinga non-volant mammal assemblages.** Relationship between differences in species richness (ΔS) and geometric mean $\log_{10}$(body mass) in the future scenarios (a) SSP245 – 2060, (b) SSP245 – 2100, (c) SSP585 – 2060, (d) SSP585 – 2100. The four levels of extrapolation constraints – MOP00, MOP70, MOP80, and MOP90 – respectively indicate species-level projections including regions with at least 00%, 70%, 80%, and 90% of environmental similarity with the training data. Symbol colours indicate the current species richness (darker = higher richness) per 10 × 10 km cell. Symbol size is proportional to the current spatial beta-diversity ($\beta_{SIM}$). Pearson correlations at the top of each panel were based on spatially corrected degrees of freedom.

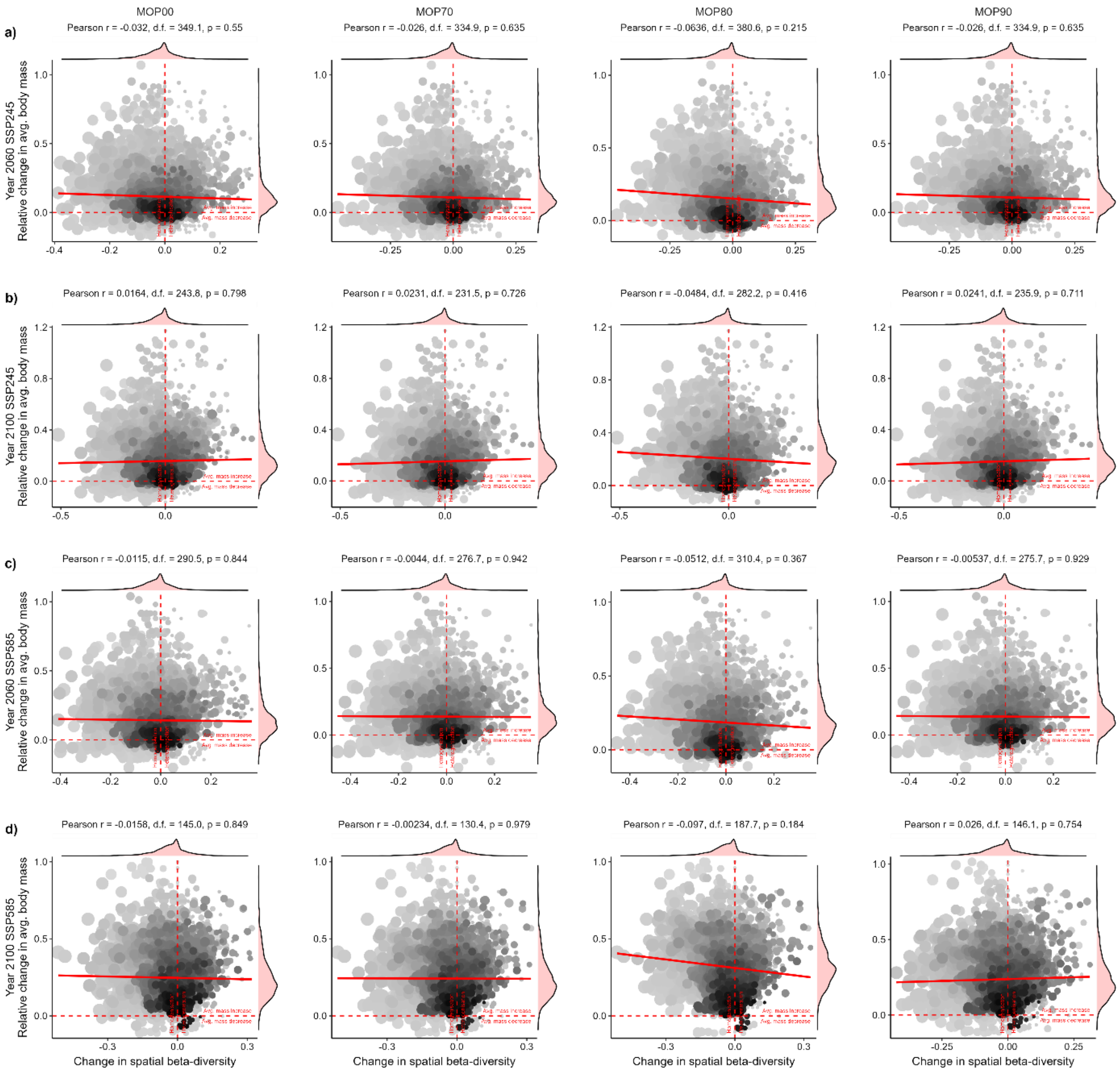


**Fig. S15. Relationship between change in spatial beta-diversity and average body mass in Caatinga non-volant mammal assemblages.** Relationship between change in beta-diversity ($\Delta\beta_{SIM}$) and geometric mean $\log_{10}$ (body mass) in the future scenarios (a) SSP245 – 2060, (b) SSP245 – 2100, (c) SSP585 – 2060, (d) SSP585 – 2100. The four levels of extrapolation constraints – MOP00, MOP70, MOP80, and MOP90 – respectively indicate species-level projections including regions with at least 00%, 70%, 80%, and 90% of environmental similarity with the training data. Symbol colours indicate the current species richness (darker = higher richness) per 10 × 10 km cell. Symbol size is proportional to the current spatial beta-diversity ($\beta_{SIM}$). Pearson correlations at the top of each panel were based on spatially corrected degrees of freedom.

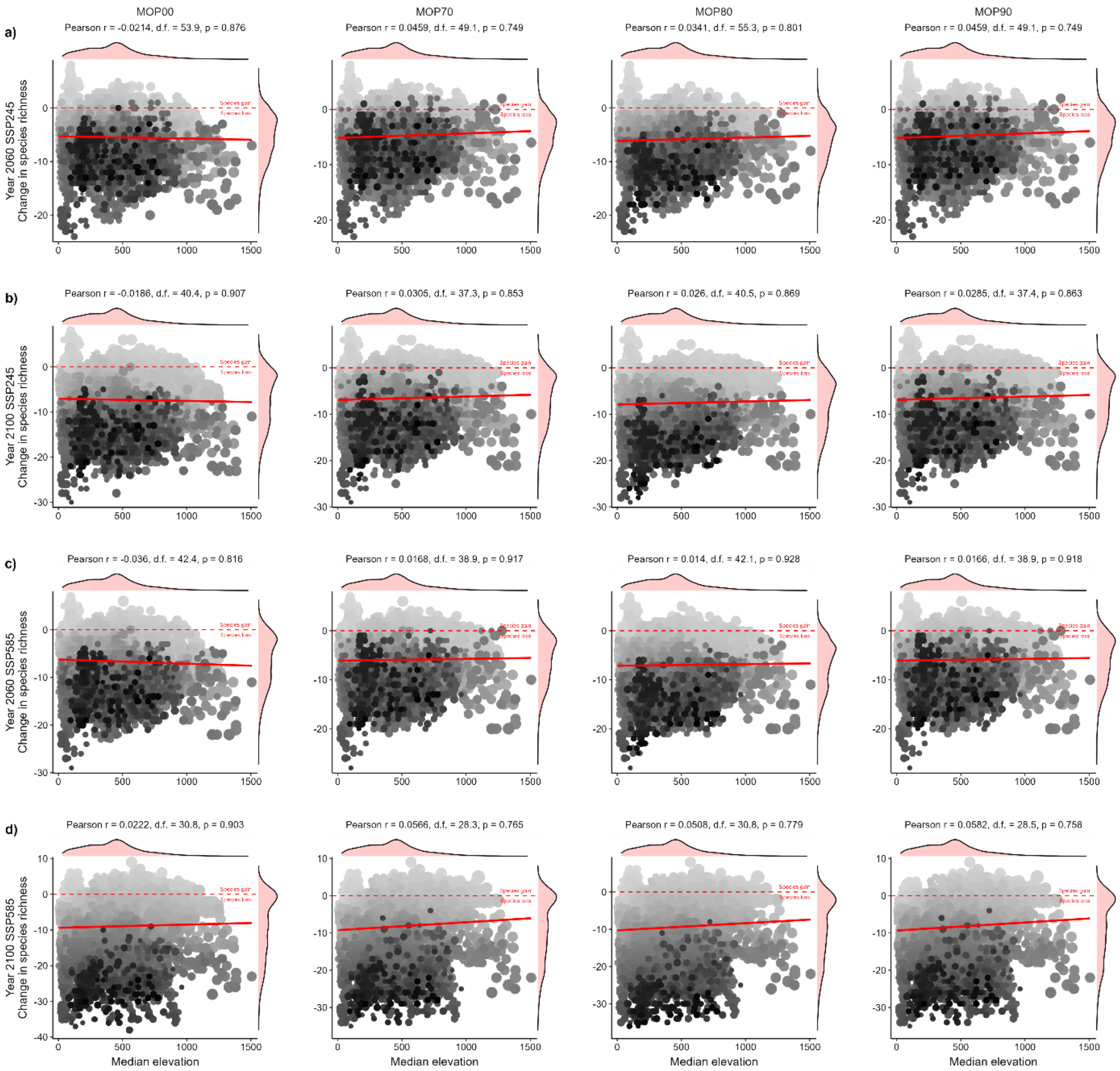


**Fig. S16. Relationship between change in species richness and elevation in Caatinga non-volant mammal assemblages.** Relationship between differences in species richness (ΔS) and within-cell median elevation in the future scenarios (a) SSP245 – 2060, (b) SSP245 – 2100, (c) SSP585 – 2060, (d) SSP585 – 2100. The four levels of extrapolation constraints – MOP00, MOP70, MOP80, and MOP90 – respectively indicate species-level projections including regions with at least 00%, 70%, 80%, and 90% of environmental similarity with the training data. Symbol colours indicate the current species richness (darker = higher richness) per 10 × 10 km cell. Symbol size is proportional to the current spatial beta-diversity ($\beta_{SIM}$). Pearson correlations at the top of each panel were based on spatially corrected degrees of freedom.

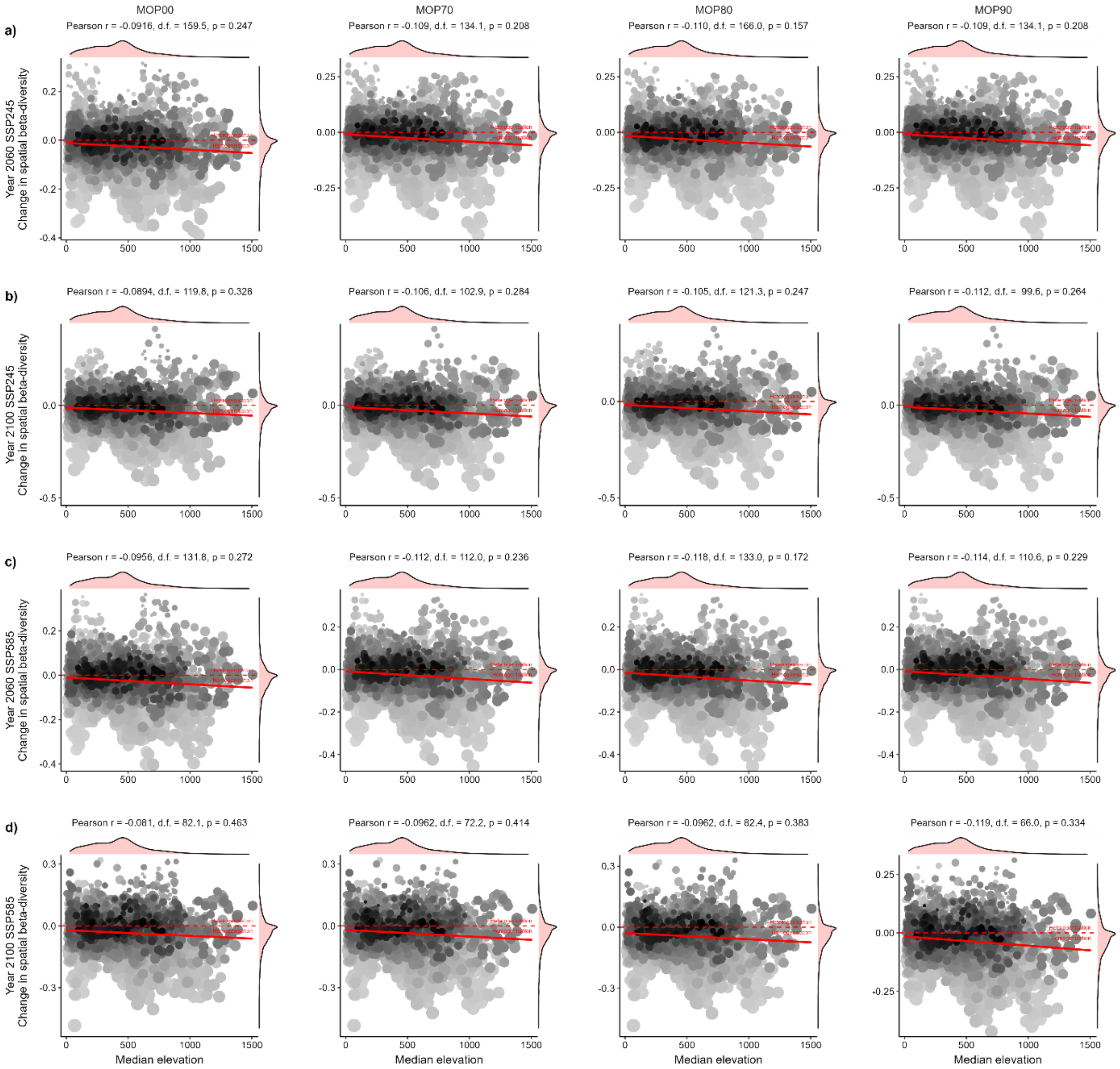


**Fig. S17. Relationship between change in spatial beta-diversity and elevation in Caatinga non-volant mammal assemblages.** Relationship between change in beta-diversity ($\Delta\beta_{SIM}$) and within-cell median elevation in the future scenarios (a) SSP245 – 2060, (b) SSP245 – 2100, (c) SSP585 – 2060, (d) SSP585 – 2100. The four levels of extrapolation constraints – MOP00, MOP70, MOP80, and MOP90 – respectively indicate species-level projections including regions with at least 00%, 70%, 80%, and 90% of environmental similarity with the training data. Symbol colours indicate the current species richness (darker = higher richness) per 10 × 10 km cell. Symbol size is proportional to the current spatial beta-diversity ($\beta_{SIM}$). Pearson correlations at the top of each panel were based on spatially corrected degrees of freedom.

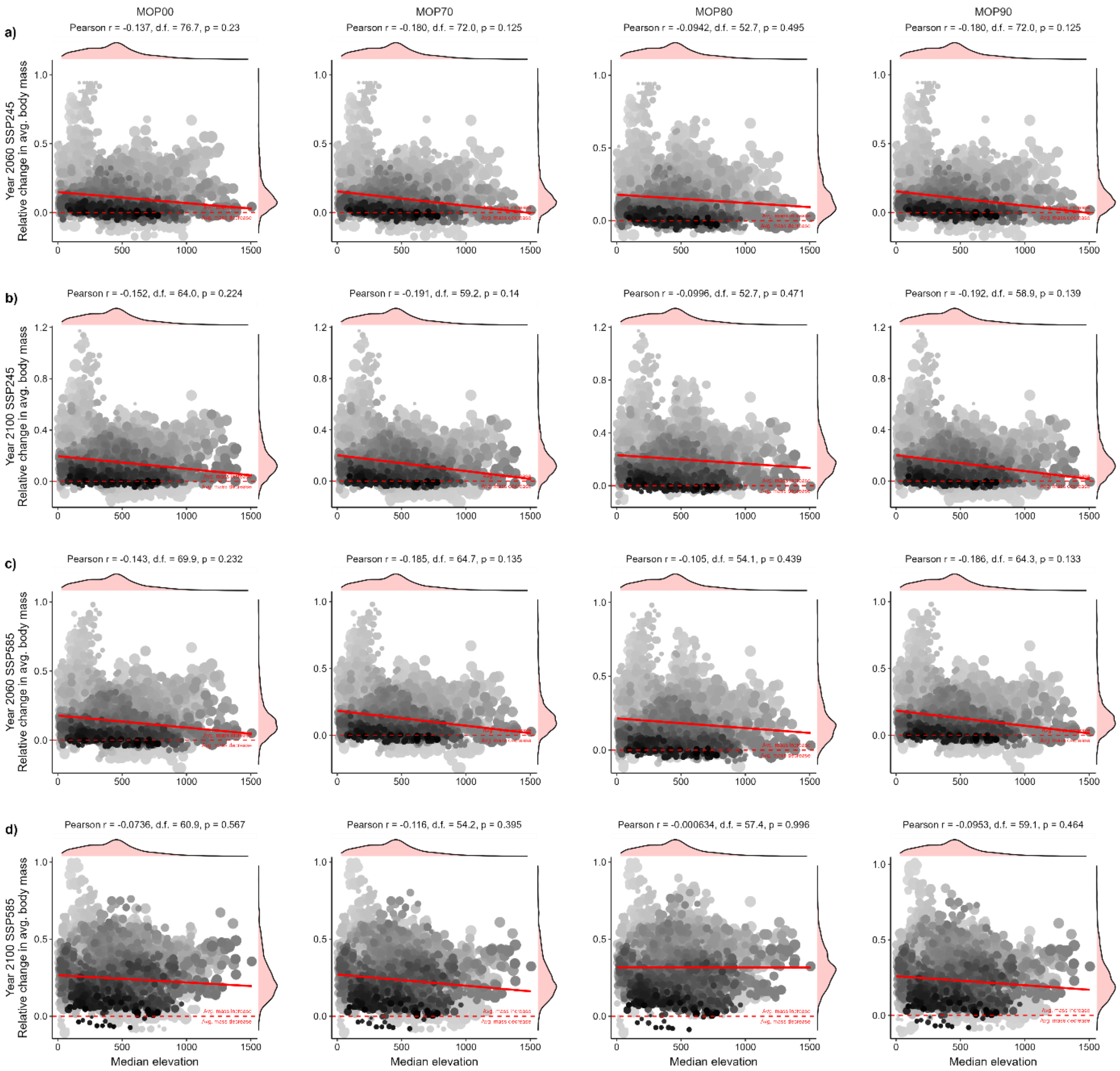


**Fig. S18. Relationship between relative change in average body mass and elevation in Caatinga non-volant mammal assemblages.** Relationship between differences relative change in geometric mean $\log_{10}$ (body mass) and within-cell median elevation in the future scenarios (a) SSP245 – 2060, (b) SSP245 – 2100, (c) SSP585 – 2060, (d) SSP585 – 2100. The four levels of extrapolation constraints – MOP00, MOP70, MOP80, and MOP90 – respectively indicate species-level projections including regions with at least 00%, 70%, 80%, and 90% of environmental similarity with the training data. Symbol colours indicate the current species richness (darker = higher richness) per 10 × 10 km cell. Symbol size is proportional to the current spatial beta-diversity ($\beta_{SIM}$). Pearson correlations at the top of each panel were based on spatially corrected degrees of freedom.

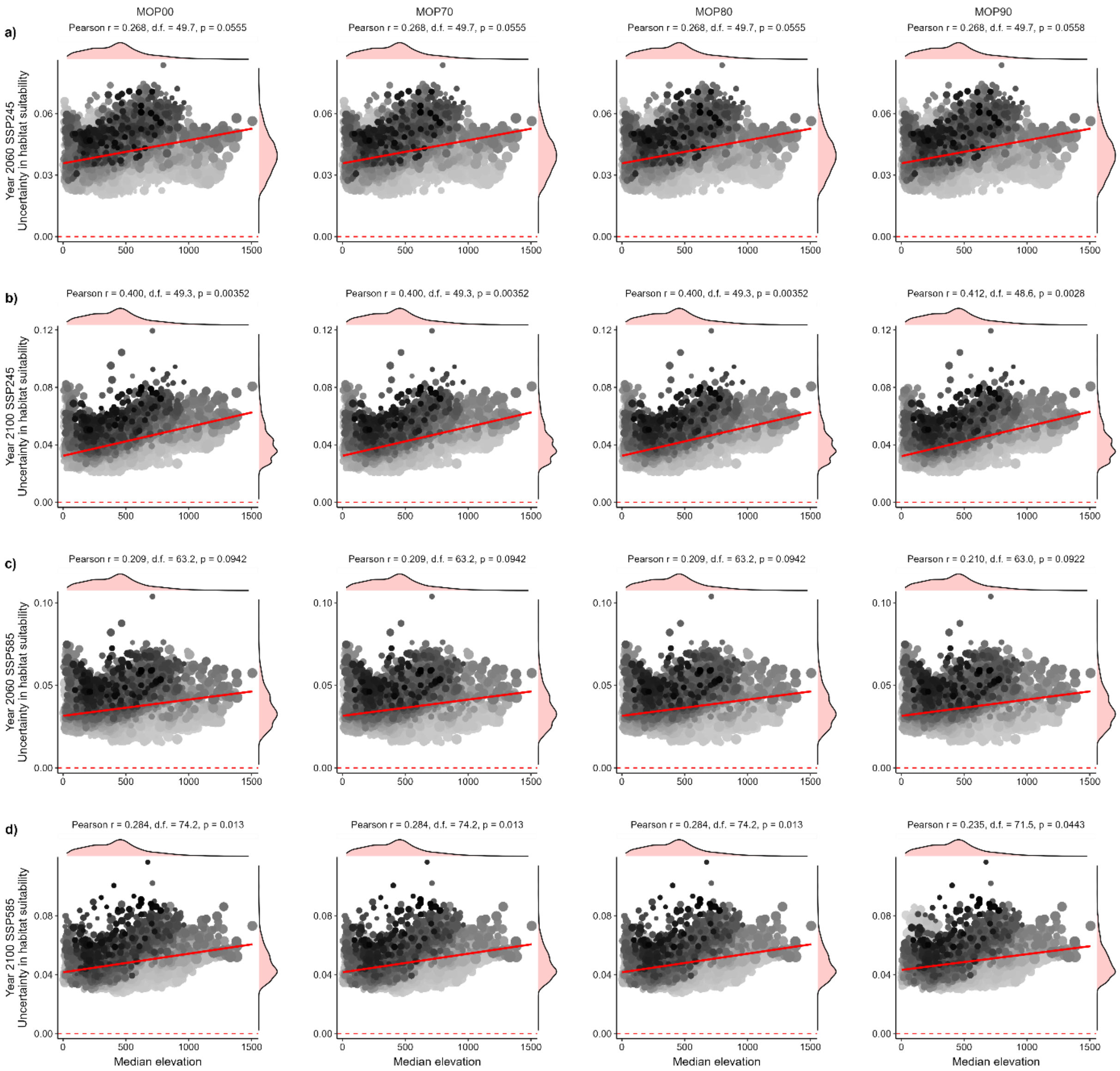


**Fig. S19. Relationship between change in model uncertainty propagation and elevation in Caatinga non-volant mammal assemblages.** Relationship between aggregated model uncertainty and within-cell median elevation in the future scenarios (a) SSP245 – 2060, (b) SSP245 – 2100, (c) SSP585 – 2060, (d) SSP585 – 2100. The four levels of extrapolation constraints – MOP00, MOP70, MOP80, and MOP90 – respectively indicate species-level projections including regions with at least 00%, 70%, 80%, and 90% of environmental similarity with the training data. Symbol colours indicate the current species richness (darker = higher richness) per 10 × 10 km cell. Symbol size is proportional to the current spatial beta-diversity ($\beta_{SIM}$). Pearson correlations at the top of each panel were based on spatially corrected degrees of freedom.

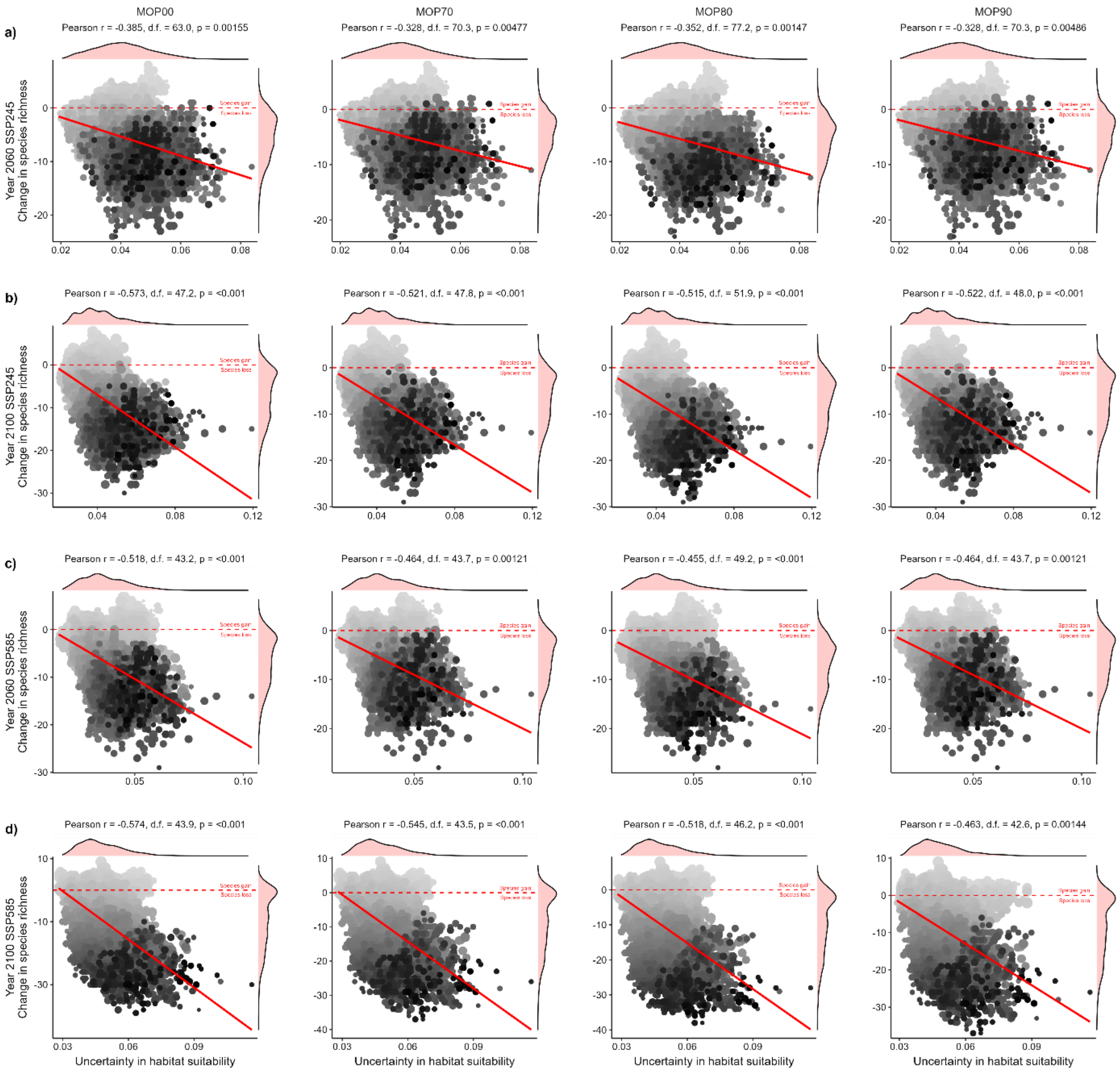


**Fig. S20. Relationship between change in species richness and model uncertainty propagation for non-volant mammal in Caatinga.** Relationship between change in species richness and aggregated model uncertainty in the future scenarios (a) SSP245 – 2060, (b) SSP245 – 2100, (c) SSP585 – 2060, (d) SSP585 – 2100. The four levels of extrapolation constraints – MOP00, MOP70, MOP80, and MOP90 – respectively indicate species-level projections including regions with at least 00%, 70%, 80%, and 90% of environmental similarity with the training data. Symbol colours indicate the current species richness (darker = higher richness) per 10 × 10 km cell. Symbol size is proportional to the current spatial beta-diversity ($\beta_{SIM}$). Pearson correlations at the top of each panel were based on spatially corrected degrees of freedom.

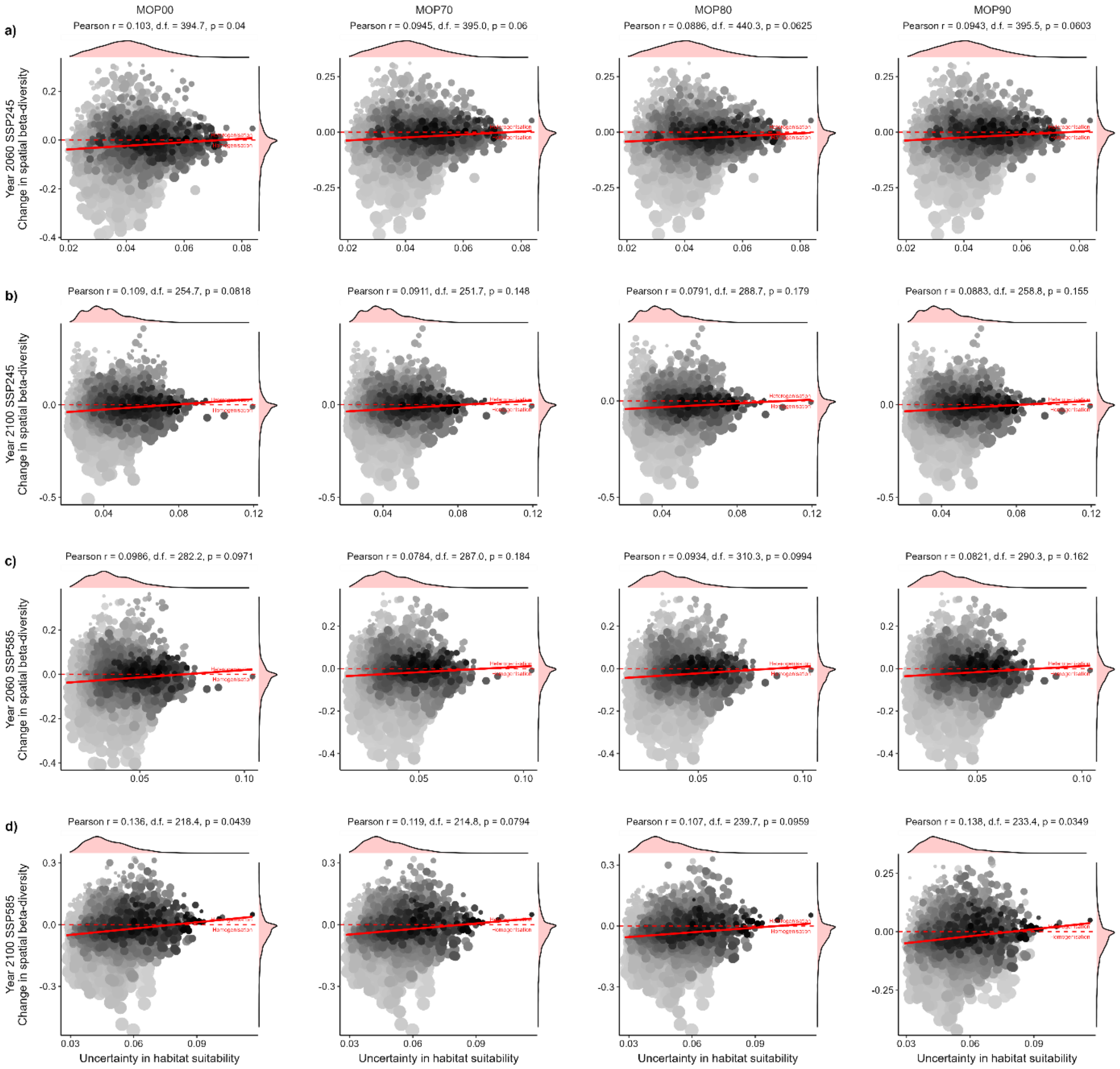


**Fig. S21. Relationship between change in spatial-beta diversity and model uncertainty propagation for non-volant mammal in Caatinga.** Relationship between change in spatial beta-diversity and aggregated model uncertainty in the future scenarios (a) SSP245 – 2060, (b) SSP245 – 2100, (c) SSP585 – 2060, (d) SSP585 – 2100. The four levels of extrapolation constraints – MOP00, MOP70, MOP80, and MOP90 – respectively indicate species-level projections including regions with at least 00%, 70%, 80%, and 90% of environmental similarity with the training data. Symbol colours indicate the current species richness (darker = higher richness) per 10 × 10 km cell. Symbol size is proportional to the current spatial beta-diversity ($\beta_{SIM}$). Pearson correlations at the top of each panel were based on spatially corrected degrees of freedom.

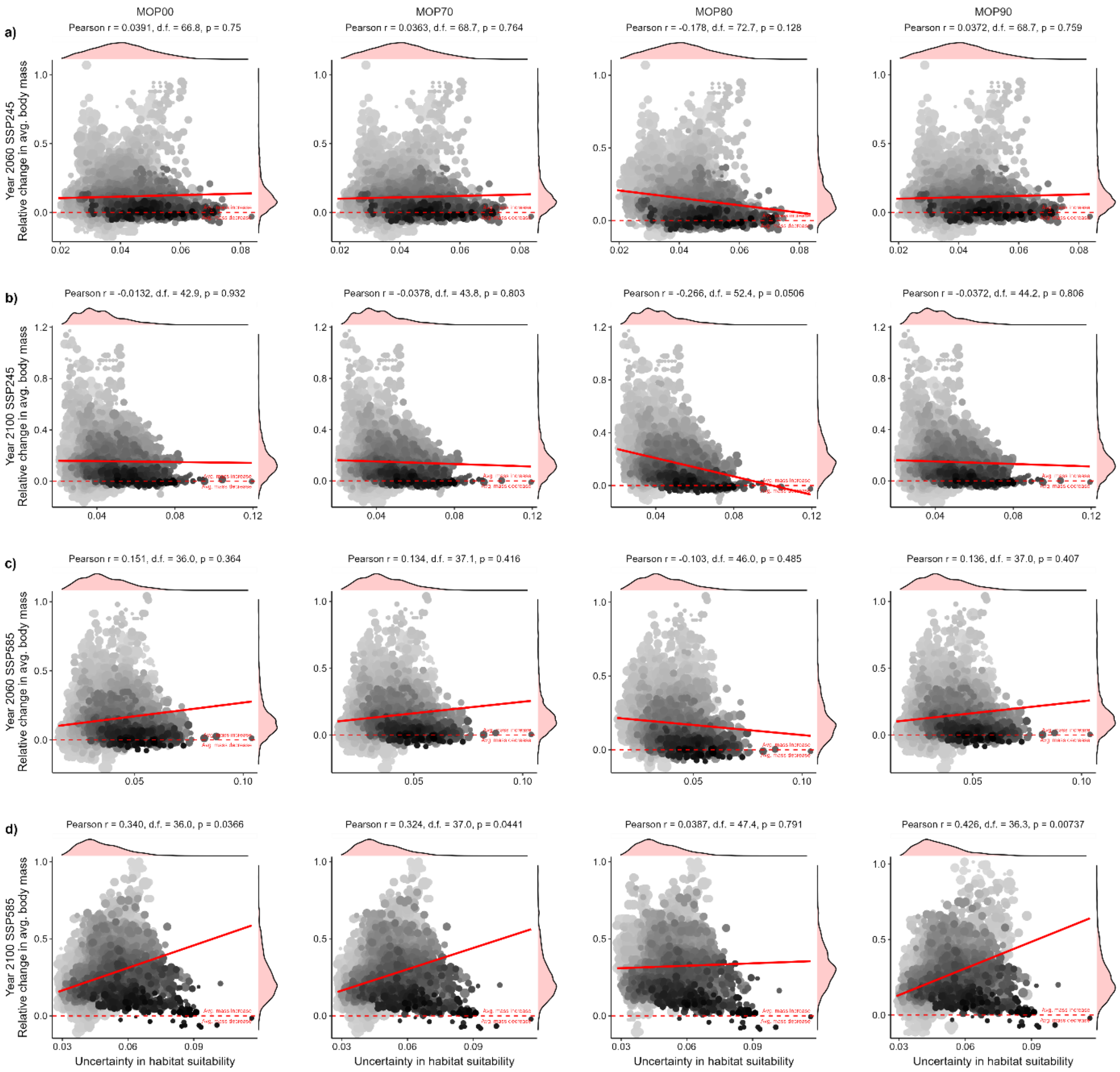


**Fig. S22. Relationship between relative change in average body mass and model uncertainty propagation for non-volant mammal in Caatinga.** Relationship between the relative change in geometric mean $\log_{10}$(body mass) and aggregated model uncertainty in the future scenarios (a) SSP245 – 2060, (b) SSP245 – 2100, (c) SSP585 – 2060, (d) SSP585 – 2100. The four levels of extrapolation constraints – MOP00, MOP70, MOP80, and MOP90 – respectively indicate species-level projections including regions with at least 00%, 70%, 80%, and 90% of environmental similarity with the training data. Symbol colours indicate the current species richness (darker = higher richness) per 10 × 10 km cell. Symbol size is proportional to the current spatial beta-diversity ($\beta_{SIM}$). Pearson correlations at the top of each panel were based on spatially corrected degrees of freedom.
